# Molecular insights into the oligosaccharide binding, recognition and hydrolysis by a fungal exo-β-(1,3)-glucanase

**DOI:** 10.1101/2024.06.05.597502

**Authors:** Barnava Banerjee, Chinmay K. Kamale, Abhishek B. Suryawanshi, Subrata Dasgupta, Santosh Noronha, Prasenjit Bhaumik

## Abstract

Carbohydrate hydrolysing enzymes assume special industrial and commercial interest as a source for yielding fermentable glucose especially for the biofuel industry. Among these enzymes, the exo-β-(1,3) glucanases are promising for industrial use as they hydrolyze sugars such as laminarin, a major constituent of the algal cell wall. Exploring the structure and function of these enzymes is of particular interest for the improvement of their functional properties for industrial use. We report the structural and biochemical characterizations of *Aspergillus oryzae* exo-β-(1,3) glucanase (AoBgl). We have expressed, purified, and performed biochemical characterizations of the recombinant AoBgl. Purified AoBgl is found to hydrolyse β-(1,3)-glycosidic linkages present in the oligosaccharide laminaritriose and the polysaccharide, laminarin effectively while retaining >50% activity at glucose concentrations of around 1.5M. We have determined three high-resolution structures of AoBgl: (a) apo form at 1.75 Å, (b) complexed form with bound disaccharide at 1.73 Å and (c) glucose-bound form at 1.20 Å. Sequence analysis and structural comparison indicate that AoBgl belongs to the GH5 sugar hydrolase family. The sugar-bound structures reveal the mode of substrate binding and interactions at the active site of AoBgl. Further, molecular dynamics simulation and mutational studies indicate that AoBgl can effectively bind trisaccharides and higher oligosaccharides. Our biochemical and structural data provide detailed molecular insights into the active site of this GH5 enzyme and would be helpful in the rational engineering of glycosyl hydrolases belonging to similar families for industrial use.

## Introduction

The hydrolysis of plant cell wall polysaccharides is of industrial and commercial interest in the biofuel industry as biofuels are clean burning and renewable energy resources (1). The increase in human population over recent years is coupled with a tremendous increase in energy requirements. This has put massive stress on the reserves of fossil fuels, which are the most commonly used forms of energy. The depleting reserves of fossil fuels as well as the detrimental effects towards the environment due to their combustion have led towards an increase in emphasis towards developing alternative sources of energy (2). Biofuels, which mainly includes bioethanol, is one such promising energy alternative. In recent years, the global interest has shifted tremendously towards the application of biofuels for meeting the energy requirements of the world population. All the G20 countries are committed to increasing the use of biofuels to meet their energy requirements (3). This means that there is a need for substantial research to explore various avenues of obtaining bio-fuels. One of the most important aspects is the source of such fuels. Most commonly, it is obtained from the saccharification of complex polysaccharides of plant origin to produce fermentable glucose, which is then converted to ethanol by fermentation. (4) The cellulosic biomass obtained from terrestrial plant cell walls has been explored extensively for biofuel production. Cellulosic biomass derived from plant cell walls include a particularly recalcitrant compound, lignin which requires extensive pre-treatment with caustic acid or alkali for its removal (5) (6). Also, some of these plants are cultivated as a source of food or fodder. Hence, constant competition exists for their use in agriculture and animal husbandry.

In contrast, algal biomass is an emerging and very promising source of raw materials for bioethanol production (7) as they do not contain recalcitrant compounds such as lignin and they can be more effectively bio-converted into simple sugars. Further, brown algae such as *Laminaria pallida* can produce biomass having a wet weight of around 14kg/m2 of the kelp bed, which is higher than the productivity of terrestrial plants like sugarcane, which can produce biomass of around 6.1-9.5 kg/m^2^/year (8). However, the hydrolysis of cell wall components of algal sources requires enzymes which are distinct from those which hydrolyse cellulosic materials. One of the most abundant biopolymers present in the cell wall of brown algae is laminarin (9) and it constitutes almost 35% of algal dry weight. Laminarin is a polymer of β-D glucose linked via β-1,3-glucan linkages (10). Laminarin is hydrolysed into glucose by enzymes known as laminarinases or exo-β-(1,3) glucanases (8). These enzymes cleave off glucose units from the reducing end of the polymer laminarin. Enzymatic hydrolysis of laminarin from brown algae has been indicated as a promising source of obtaining fermentable glucose in recent years (12). The enzymes that hydrolyze β-1,3-glucan linkage are present in several terrestrial fungal species such as those belonging to the *Aspergillus* genus (13, 14). These enzymes are crucial for the breakdown of cell wall components of such fungi during the process of hyphal branching and conidial germination (15). Most of these enzymes are secreted into the environment and can be exploited for industrial applications such as third-generation bioethanol production from algal biomass (16).

Exo-β-(1,3)-glucanases (EC 3.2.1.58) belong to a family of enzymes called glycosyl hydrolases (GHs). The Carbohydrate Active enZymes (CAZy) (http://www.cazy.org) is a comprehensive database that stores information regarding various families of GH enzymes based on their sequences. According to CAZy, exo-β-(1,3)-glucanases belong to several families of Glycosyl Hydrolases such as GH3, GH5, GH17 and GH55 (17). Among these, GH5 family of glycosyl hydrolase enzymes assumes particular interest as this family of enzymes includes both endoglucanases which hydrolyse as polysaccharide chain internally as well as exoglucanases, which hydrolyse the polysaccharide chain from its ends (18–20). GH5 enzymes have a TIM barrel structural fold. Interestingly, GH1 family of glycosyl hydrolase enzymes which include β-glucosidases (20–22) also have the same structural architecture. β-glucosidases are the enzymes which hydrolyse the β-(1,4) glycosidic bond in disaccharide cellobiose, to yield glucose. The enzymes belonging to the GH1 families have been extensively studied both structurally and biochemically. However, the studies on their GH5 counterparts are relatively few. GH5 enzymes have enormous flexibility with respect to the type of substrate they can hydrolyse. As GH5 enzymes can hydrolyse both β-(1,4) as well as β-(1,3) linkages in sugars, such functional properties make them diverse with respect to their applications (23–27). There have been several reports on the utility of exo-β-(1,3)-glucanases towards various industrial and agricultural applications (28). However, the lack of high-resolution structural information has limited the efforts to engineer these proteins for effective industrial use.

In this study, we report, for the first time, recombinant expression, purification, and characterizations of a GH5 exo-β-(1,3) glucanase enzyme from *Aspergillus oryzae* (AoBgl) from a bacterial expression system. Pure recombinant AoBgl was found to hydrolyse laminarin effectively at a temperature of 50°C and pH of 5.5. We solved high-resolution crystal structures of AoBgl in apo, cellobiose and glucose-bound forms, which revealed the mode of substrate binding and the dynamics of binding of sugar moieties inside the active site crater. Our structural and biochemical data would aid in the development of superior variants of AoBgl for industrial biofuel production.

## Results

### Expression and purification of AoBgl

AoBgl was expressed successfully as a soluble recombinant protein in *E. coli* BL21(DE3) cells. The enzyme was purified by three cycles of chromatographic techniques; beginning with Ni-NTA affinity chromatography, followed by anion-exchange chromatography and finally with size exclusion chromatography. The initial trials with the Ni-NTA chromatography purification did not yield a substantial amount of AoBgl in pure form as most of the protein was eluted with imidazole concentrations of 25-50 mM, which also co-eluted several contaminant proteins. Performing size exclusion chromatography with the concentrated fractions of Ni-NTA failed to remove all such contaminants. Hence, there was a requirement for another intermediate purification step, which was, in this case, anion exchange chromatography. The theoretical isoelectric point (pI) of AoBgl is 4.2, which was in the acidic range. We hypothesized that dilution of the eluted protein samples from Ni-NTA containing AoBgl in an appropriate buffer of pH 5.2 would cause precipitation of most of the bacterial proteins as their isoelectric points are near pH of 5.0. Bacterial proteins at this pH would be devoid of a net surface charge, thereby reducing their solubility in an aqueous solvent and aiding in their precipitation. But AoBgl, owing to its low pI would still be negatively charged at pH 5.2, remain in solution and be purified by an anion exchanger. Hence, an anion exchange chromatography step was carried out; and significantly pure AoBgl could be obtained with the elution using 250 mM sodium chloride. Size exclusion chromatography was used as a final step removing any other contaminants. Size exclusion chromatography elution profile showed a single peak corresponding to the monomeric molecular weight of the enzyme thereby confirming that highly pure AoBgl was obtained (Figure S1). Our study reports the first successful recombinant expression and purification of AoBgl from a bacterial expression system.

### AoBgl is a GH5 enzyme: Biochemical characterizations and bioinformatics analysis

The biochemical characterizations of AoBgl were initially performed with p-NPG hydrolysis assays. An earlier study of this enzyme purified from the native source reported AoBgl as a highly glucose tolerant β-glucosidase with a capacity of hydrolysing β-(1,4)-glycosidic bonds. (29). The artificial substrate p-NPG was used for characterizations of AoBgl extracted from a native source, and the enzyme was reported to be active on cellobiose. Our biochemical assays (discussed in the following sections) with the pure recombinant enzyme showed that AoBgl could effectively carry out p-NPG hydrolysis but, surprisingly, was inactive against cellobiose, which is the real substrate for the β-glucosidases. On comparing the sequence of AoBgl, it was found that it aligned well with enzymes belonging to exo-β-(1,3)-glucanases, which carry out hydrolysis of glucose polymers linked with β-(1,3) glycosidic linkages such as laminarin. Sequence analysis of AoBgl using Pfam server classified AoBgl in family 5 of glycosyl hydrolases (GH5). Sequence alignments also showed that AoBgl aligned better with the GH5 family enzymes than GH1 family β-glucosidases with respect to its catalytic motifs (Figure 1, S2 and S3). Furthermore, β-glucosidases are characterized by two conserved motifs, NEP and ENG (30–33), where each motif hosts the catalytic glutamates. AoBgl was found to lack the ENG motif which was another evidence of it not being a β-glucosidase.

**Figure 1:**
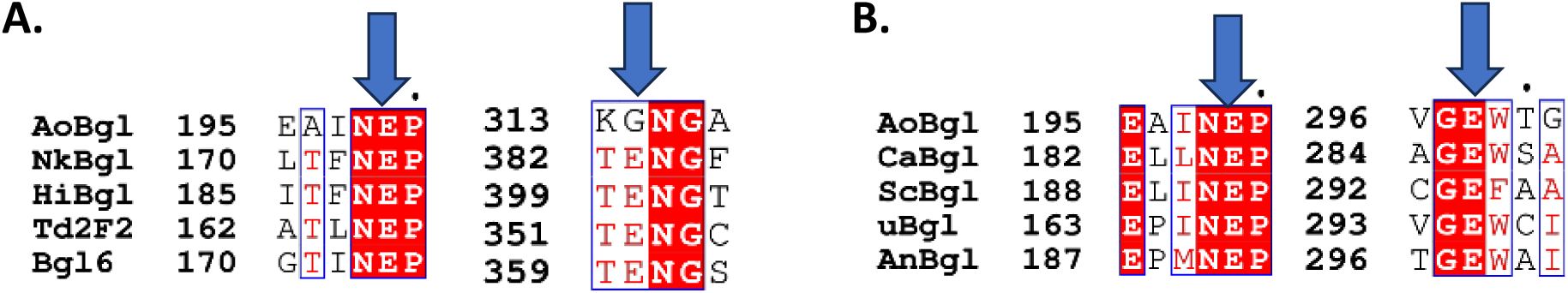
Sequence analysis of AoBgl. (A) Comparison of AoBgl sequence with characterized β-glucosidases belonging to GH1 family. For this alignment β-glucosidases from *Neotermes koshunensis* (NkBgl), *Humicola insolens* (HiBgl), and soil metagenome (Td2F2 and Bgl6) have been considered. (B) Comparison of AoBgl sequence with GH5 family enzymes. For this alignment GH5 exo-β-(1,3)-glucanases from *Candida albicans* (CaBgl), *Saccharomyces cerevisiae* (ScBgl), uncultured bacterium (uBgl) and *Aspergillus niger* (AnBgl) have been considered. The motifs hosting the catalytic glutamate residue are conserved in alignment (B), whereas one of the catalytic motifs (ENG) is absent in (A). The positions of the catalytic glutamate residues in each motif have been indicated with a blue arrow.

The hydrolysis activity of purified AoBgl was investigated on β-(1,3) linked sugars such as the trisaccharide, laminaritriose and the polysaccharide, laminarin. The enzyme hydrolysed these sugars successfully to release glucose. However, AoBgl showed extremely low activity on β-(1,4) linked oligosaccharides such as cellobiose and cellotriose. This indicates that AoBgl preferentially hydrolyses β-(1,3) linked oligosaccharides whereas there was no detectable activity on their β-(1,4) linked counterparts. The relative activities of AoBgl on a variety of sugar substrates have been shown in Table 2 and Figure 2F.

**Figure 2:**
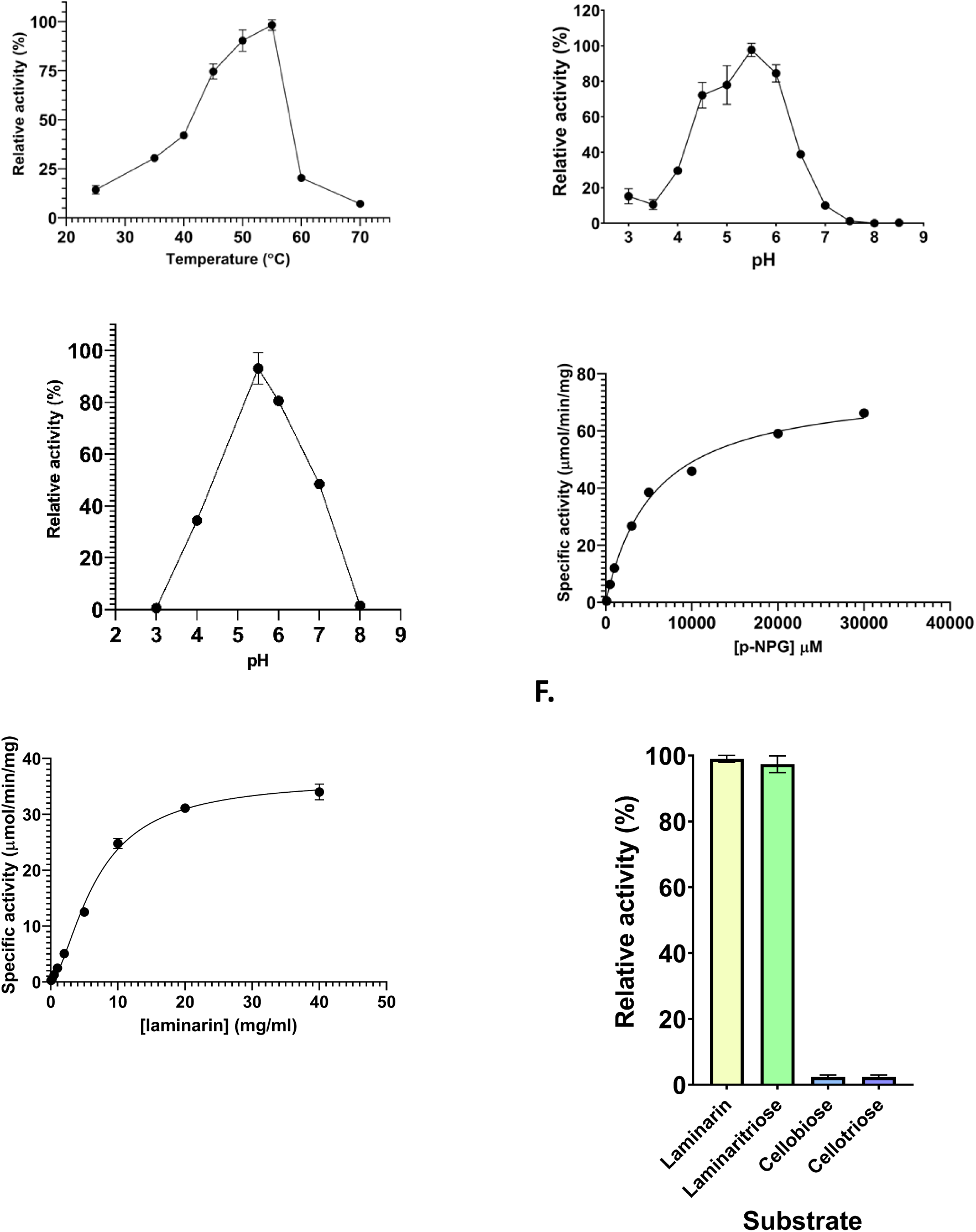
Biochemical properties of AoBgl. (A) Optimum temperature of p-NPG hydrolysis. (B) Optimum pH for p-NPG hydrolysis. (C) Optimum pH for laminarin hydrolysis. (D) Kinetic characterizations of AoBgl using p-NPG hydrolysis assays. (E) Kinetic characterizations of AoBgl using laminarin hydrolysis assays. (F) Relative activities of AoBgl on different sugar substrates. Error bars show standard errors in reading obtained in triplicates.

#### Temperature optima and pH optima of AoBgl

The optimum temperature for p-NPG hydrolysis of AoBgl is 55°C. However, the enzyme also shows almost 80% of activity at 50°C (Figure 2A). Following its optimum temperature, there was a steep decline in activity with an increase in temperature. The optimum pH for AoBgl activity was determined by performing the assays based on p-NPG hydrolysis. The activity versus pH profile (Figure 2B) of AoBgl was a bell-shaped curve indicating the enzyme’s maximum activity at pH 5.5. However, AoBgl can retain more than 80% of its activity at pH 4.5-5.0, indicating this enzyme’s suitability for industrial applications. The temperature and pH profiles of AoBgl make it a promising candidate for the industrial saccharification process. The pH profile of AoBgl was very different when the assays were carried out using laminarin as the substrate; the curve was very steep; following optimal laminarin hydrolysis at pH 5.5, there was a steep decline in enzyme activity (Figure 2C). These observations indicate that the p-NPG-based assays for biochemical characterizations of glycosyl hydrolases might not represent the actual activity of these enzymes as it should be towards their natural substrates.

#### Kinetic parameters of AoBgl

Kinetic parameters of AoBgl were determined using p-NPG hydrolysis assays (Figure 2D). AoBgl showed Michaelian enzyme kinetics with K_M_ of 5.78mM and a V_max_ of 78.088 µmol/min/mg. AoBgl kinetics performed using laminarin revealed an interesting and unique property of the enzyme. AoBgl showed slight allosteric behaviour towards the substrate laminarin. The Hill coefficient was calculated to be around 1.6, indicating a slightly positive allosteric nature of the enzyme (Figure 2E). The kinetics of cellobiose hydrolysis was also attempted to be performed as AoBgl purified from a native source was reported to hydrolyse this disaccharide. However, we found out that AoBgl does not hydrolyse cellobiose under standard assay conditions. This result confirms that AoBgl cannot hydrolyse β-(1,4) glycosidic bonds present in cellobiose.

#### Product inhibition of AoBgl

The hydrolysis reaction of p-NPG was carried out by AoBgl in the presence of increasing concentrations of glucose (Figure 3). After an initial increase in the activity up to 0.5 mM glucose, a gradual decrease in activity was seen with an increase in glucose concentration. This increase in activity with the addition of glucose can be due to a phenomenon called glucose stimulation as reported for many glycosyl hydrolases belonging to the GH1 family (34, 35). This happens when glucose molecules occupy the other sugar-binding sites on the protein molecule, thereby leaving only the catalytic site for the p-NPG molecule to bind. This causes an increase in the activity at lower glucose concentrations. However, at higher glucose concentrations, the glucose molecules, which are now in abundance, also start occupying the sites for substrate binding. This causes a decrease in the hydrolytic activity of the enzyme. For AoBgl, it was found that p-NPG hydrolysis was inhibited by 50% at glucose concentrations of over 1.5M (Figure 3). The industrial bio-ethanol reactors generally require enzymes which are not inhibited even at higher concentrations of glucose *i.e.* retain ~50% of their activity at glucose concentrations of around 1 M since the maximum concentration of glucose that is reported in industrial reactors is around 0.7-0.8 M (36). From our study, we can conclude that AoBgl can be a very promising candidate in industrial reactors owing to its very low levels of product inhibition.

**Figure 3:**
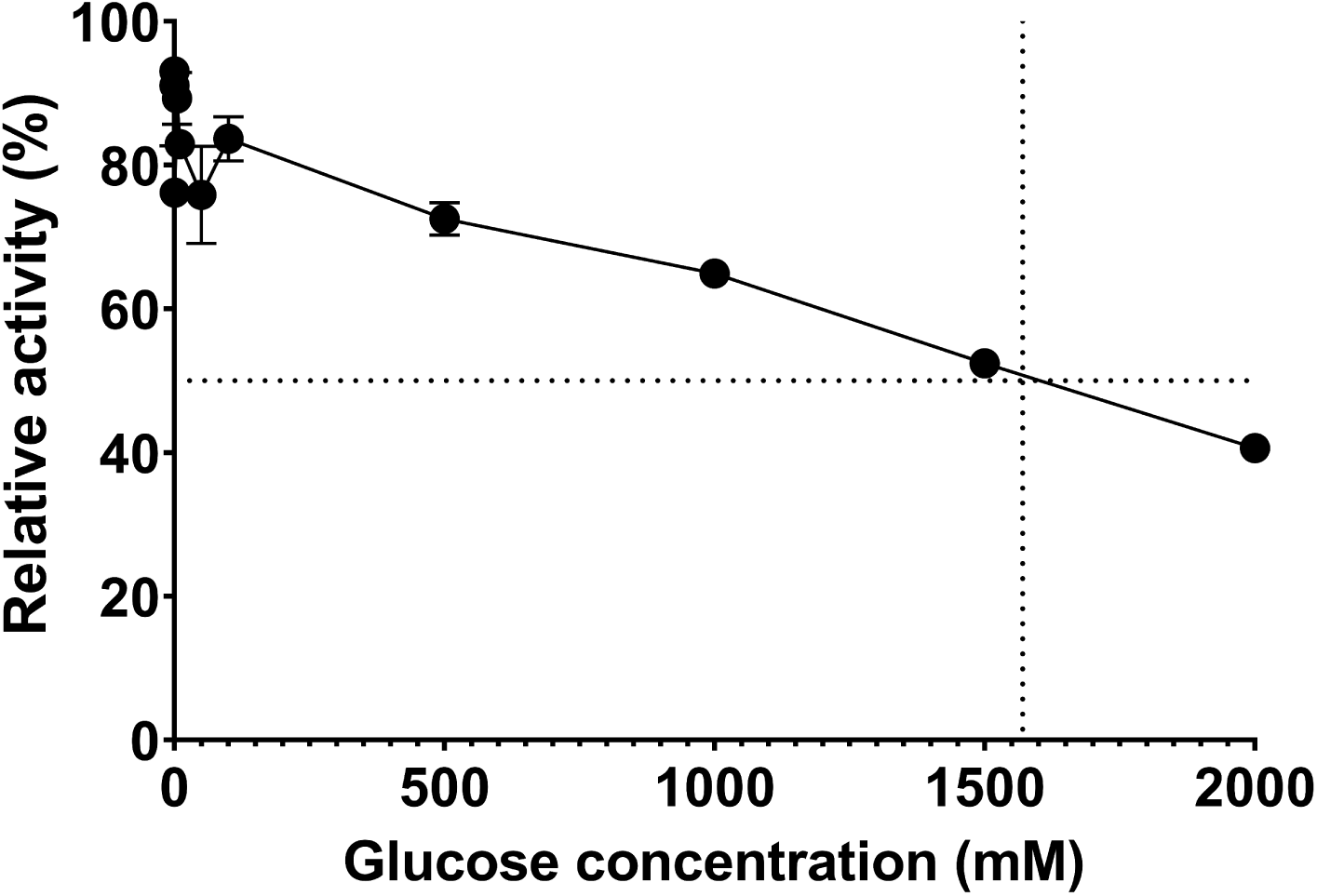
Product (glucose) inhibition in AoBgl. Changes in p-NPG hydrolytic activity of AoBgl towards p-NPG at different concentrations of product (glucose). The dotted lines indicate that AoBgl loses 50% activity at around 1.5 M glucose. Error bars show standard errors in reading obtained in triplicates.

#### Stability of AoBgl at different temperatures

One of the other major factors that determine the employability of enzymes in the industrial bio-ethanol production processes is their stability at different temperatures. Thus, the effect of temperature on the stability of AoBgl was monitored. The assays at 30°C showed that the AoBgl retained more than 80% of its activity over 48 hours of incubation at the said temperature (Figure 4). However, the enzyme lost its activity within 1 hour of incubation at its optimum temperature i.e., 55°C for catalysis.

**Figure 4:**
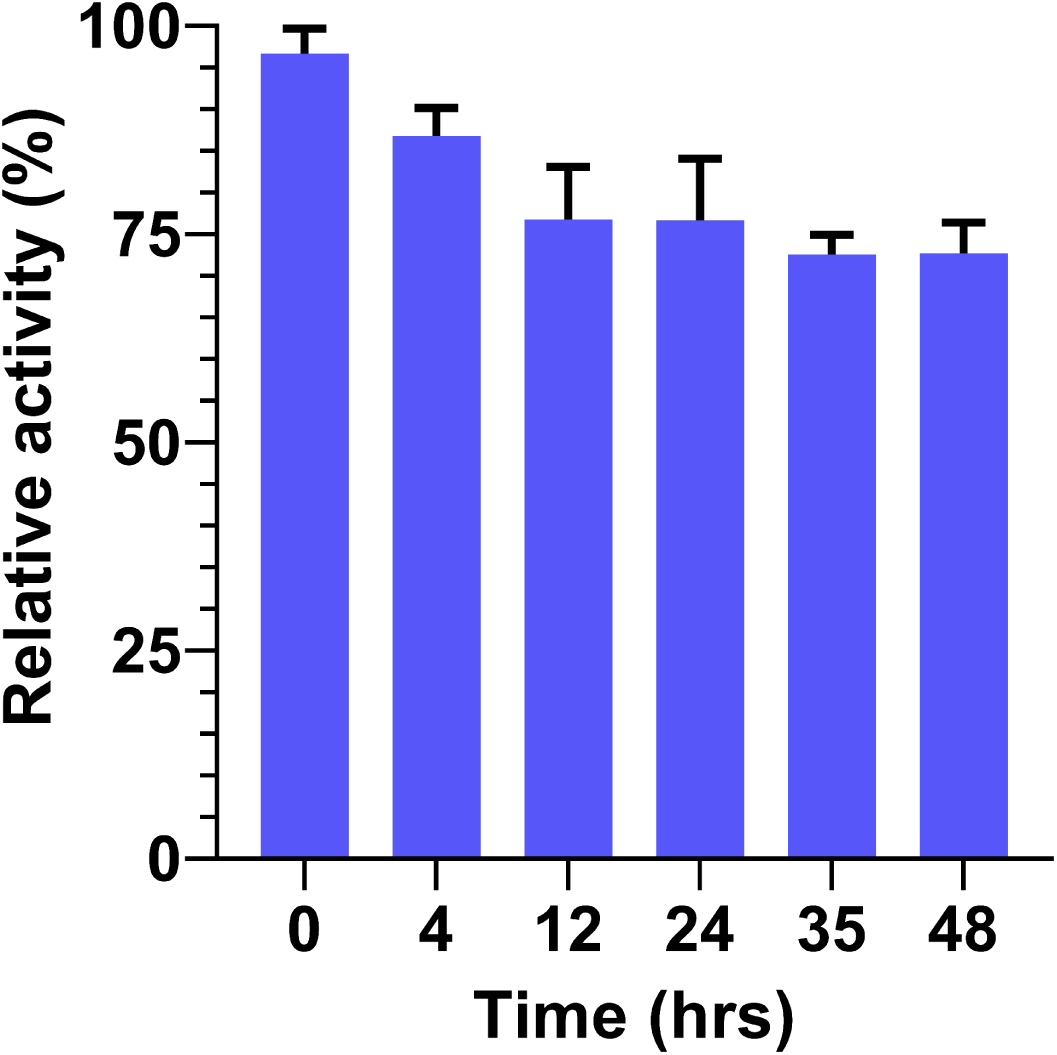
Stability of AoBgl. Relative activity of AoBgl assayed at various time intervals while incubating the enzyme at 30°C. Error bars show standard errors in reading obtained in triplicates.

The reason behind this was investigated through the temperature-mediated unfolding of AoBgl by monitoring the tryptophan fluorescence of the protein at various temperatures from 30°C-60°C. AoBgl was incubated at each temperature for one hour and readings were taken every 5 minutes. It was noted that during this incubation period, there was very little change in the fluorescence intensity at 337 nm at lower temperatures, however at higher temperatures of 55°C and 60°C, there was a steep decline in the fluorescence intensity at 337 nm which indicated unfolding of the protein at these temperatures thereby exposing the buried tryptophan residues towards the aqueous environment. The unfolding of AoBgl at these temperatures might be the cause of the observed loss of activity (Figure S5).

### Structural characterizations and analysis of AoBgl

We determined three high resolution crystal structures of AoBgl to understand the molecular details of the active site and substrate binding pocket of the enzyme. After incubation of the crystal trays at 18°C for 6 months, tiny crystals (0.1mm×0.1mm×0.1mm) of apo-AoBgl appeared in the wells where the protein was incubated with 1.6M sodium citrate tribasic dihydrate at pH 6.5 and 0.2 M sodium tartarate with 20% (w/v) polyethylene glycol (PEG) at pH 6.9. The mother liquor served as a cryoprotectant for the former, while 30% glycerol was used as a cryoprotectant for the latter. The apo-AoBgl crystal diffracted to a high resolution of 1.75 Å for the apo-enzyme. The high-resolution crystal structures of AoBgl as apo-enzyme and complexed with cellobiose and as well as glucose were determined. All the structures were well refined with low R-factors and good stereochemical parameters (Table 1). The structural details of AoBgl as an apo-enzyme as well as a sugar-bound complex have been presented in the sections below.

**Table 1:** Data collection and refinement statistics for Apo, disaccharide (cellobiose)-bound and glucose-bound forms of AoBgl.

|  | Apo-AoBgl | AoBgl-Cellobiose | AoBgl-Glucose |
| --- | --- | --- | --- |
| **Data collection statistics<sup>a</sup>** |  |  |  |
| Space group | *P*3<sub>2</sub>21 | *P*3<sub>2</sub>21 | *P*3<sub>2</sub>21 |
| Unit cell parameters<br>a, b, c (Å)<br>$\alpha$, $\beta$, $\gamma$ (°) | 81.49, 81.49, 115.16<br>90, 90, 120 | 80.87, 80.87, 113.64<br>90, 90, 120 | 81.29, 81.29, 114.73<br>90, 90, 120 |
| Temperature (K) | 100 | 100 | 100 |
| Wavelength (Å) | 1.5418 | 1.5418 | 0.97893 |
| Resolution range (Å) | 70.57 - 1.75 (1.78-1.75) | 38 - 1.73 (1.83-1.73) | 20-1.2 (1.4-1.2) |
| R<sub>merge</sub> (%) | 9 (34.0) | 9.5 (59.8) | 11.1 (87.5) |
| Completeness | 99.90 (98.90) | 99.9 (99.8) | 98.3 (96.6) |
| Mean I/$\sigma$ (I) | 18.10 (3.80) | 19.78 (3.87) | 13.72 (3.00) |
| Total reflections | 412868 (11124) | 469111 (63238) | 1369669 (462153) |
| Unique reflections | 45212 (2452) | 45479 (6940) | 134619 (48574) |
| Redundancy | 9.10 (4.50) | 10.31 (9.11) | 10.17 (9.51) |
| Wilson B-factor (Å<sup>2</sup>) | 8.19 | 15.75 | 11.00 |
| CC<sub>1/2</sub> (%) | 100 (86) | 99.9 (88.3) | 99.9 (85.8) |
| Number of molecules in ASU | 1 | 1 | 1 |
| **Refinement statistics** |  |  |  |
| Resolution (Å) | 70.57 - 1.75 | 35.03 - 1.73 | 19.53 - 1.20 |
| No. of reflections | 45171 | 43205 | 127482 |
| R<sub>work</sub> (%) | 14.39 | 13.96 | 15.31 |
| R<sub>free</sub> (%) | 17.18 | 16.33 | 18.17 |
| Protein atoms | 3000 | 3008 | 2885 |
| Water molecules | 316 | 247 | 585 |
| Sodium ion | 1 | 1 | 1 |
| Chloride ion | 0 | 0 | 1 |
| Cellobiose molecule | 0 | 1 | 0 |
| Glucose molecule | 0 | 2 | 2 |
| Glycerol molecule | 0 | 0 | 2 |
| **Geometry statistics** |  |  |  |
| RMSD bonds (Å) | 0.01 | 0.03 | 0.01 |
| RMSD angles (°) | 1.61 | 2.32 | 1.84 |
| **Ramachandran plot** |  |  |  |
| Favoured (%) | 96.6 | 95.25 | 95.0 |
| Allowed (%) | 3.4 | 3.96 | 4.0 |
| Outliers (%) | 0.0 | 0.79 | 1.0 |
| PDB ID | 8Z2W | 8Z2X | 8Z2Y |

<sup>a</sup> Values in parentheses correspond to the highest resolution shell.

**Table 2:** Relative hydrolysis activity of AoBgl on different sugars.

| Substrate | % hydrolysis activity | Type of linkage | Number of sugar units |
| --- | --- | --- | --- |
| Laminaritriose | 100 | β-(1,3) | 3 |
| Laminarin | ~100 | β-(1,3) | n<sup>#</sup> |
| Cellobiose | <2 | β-(1,4) | 2 |
| Celotriose | <2 | β-(1,4) | 3 |

<sup>#</sup> indicates polymer of “n” sugar units

#### Overall structure and active site of apo AoBgl

The structure of AoBgl in apo form was solved at 1.75 Å resolution. The overall structure of AoBgl shows TIM barrel or (α/β)8 fold, which is a characteristic feature of enzymes belonging to the GH5 family of glycosyl hydrolases. Superposition of the refined AoBgl model was carried out with a well-characterized exo-β-(1,3) glucanase from *Candida albicans* (PDB ID 1CZ1) (37), which bears around 45% sequence identity with AoBgl. The two structures superimposed very well with core root mean square deviation (RMSD) of 0.75Å between the two structures. The structural comparison and sequence alignment identified the two catalytic glutamates as E199 and E298. E199 was identified as the general acid/base while E298 was the catalytic nucleophile.

The two catalytic glutamates are at a distance of 5.2Å (Figure 5A), which is a characteristic feature of glycosyl hydrolases that carry out carbohydrate hydrolysis by a retention mechanism (38). The residues surrounding the two catalytic glutamates include N156, H145 and N195. These residues are polar in nature and are expected to stabilize the substrate molecule by forming polar interactions with the −OH groups of the sugar moieties.

**Figure 5:**
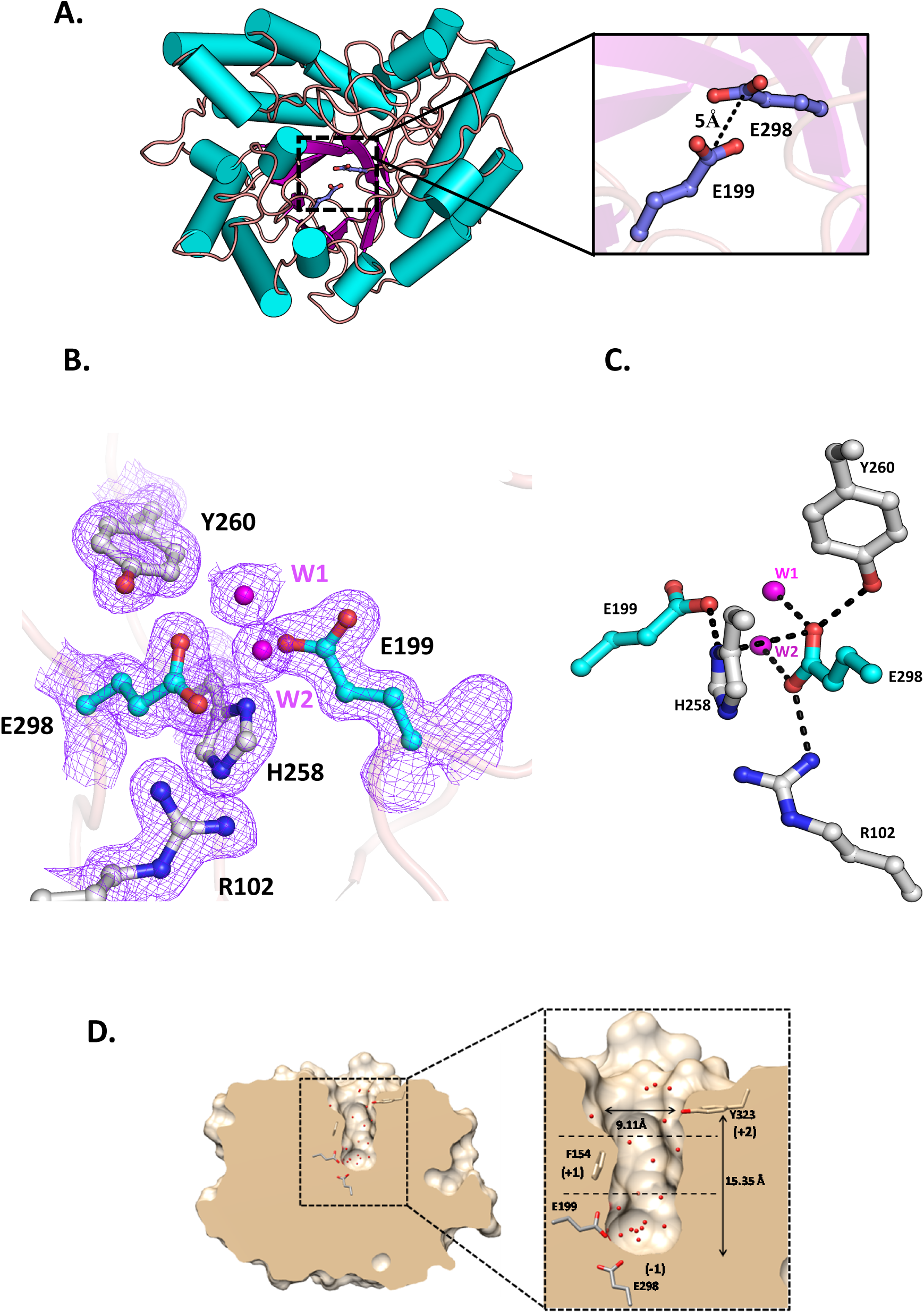
Structural features of apo-AoBgl. (A) Overall structure of AoBgl showing characteristic TIM barrel fold which is observed in GH5 enzymes. Inset: catalytic residues E199 and E298, located at a distance of 5Å, which is a characteristic feature of glycoside hydrolases showing the retaining mechanism of carbohydrate hydrolysis. (B) 2*F_o_-F_c_* map contoured at 1σ level showing the residues at the catalytic crater along with two conserved water molecules. The catalytic (E199 and E298) residues are shown in cyan and other residues in the vicinity are depicted in light grey. The water molecules are shown as magenta spheres. (C) Interactions stabilizing the catalytic crater. The catalytic glutamates (E199 and E298) and the other residues it interacts with are shown. Two water molecules shown as W1 and W2 also interact with the catalytic residues. (D) Transverse section showing the depth of the catalytic crater of AoBgl. The presence of several water molecules (shown as red spheres) at the crater is observed.

Polar residues and water molecules in the active site crater also help stabilize the catalytic glutamates by forming hydrogen bonds. We observed two water molecules (W1 and W2) which form hydrogen bonds with the catalytic nucleophile E298. The temperature factor of W1 is 6.24 Å^2^ while that of W2 (Figure 5C) is 11.65 Å^2^ indicating that these molecules are very well stabilized in the active site of AoBgl. This might be because they form strong interactions with E298. E298 is also stabilized by polar interactions with Y260 and a strong salt bridge interaction with R102. The acid/base E199 is stabilized by a single salt bridge interaction with H258, forming weak salt bridges with E298 (Figures 5B, C). The catalytic crater has an overall depth and width of 15.35 Å and 9.11 Å, respectively (Figure 5D). This deep-seated crater might be one of the factors responsible for AoBgl being highly glucose tolerant. There are several water molecules present near inside the crater which might play an important role in substrate binding and catalysis (Figure 5D).

#### Sugar binding in the active site and catalytic crater of AoBgl

The crystals were soaked in cellobiose to understand the mode of binding and interactions at the active site of the enzyme, as from our biochemical studies, we knew that AoBgl does not hydrolyze this β-(1,4) glycosidic bond. The crystal diffracted to a high resolution of 1.73 Å. Analysis of the electron density (Figure 6A) showed the presence of bound cellobiose in the crater of AoBgl. The cellobiose is bound at the entrance of the catalytic crater (Figure 6A-C). Notably, cellobiose was captured in a double conformation, which indicates that there is some amount of conformational flexibility in the manner by which cellobiose binds to the active site of AoBgl. The cellobiose molecule is stabilized by both polar and hydrophobic stacking interactions. Polar interactions involve the residues like N156 and R318 (Figure 6C). Two phenylalanine residues, F154 and F263, which also form the gate-keeper residues of the active site crater, stabilize cellobiose at the entrance of the crater by forming stacking interactions with the carbon backbone (Figure 6B). This is also consistent with what is reported in the literature about aromatic residues forming a clamp-like structure guarding the entrance of the catalytic crater (39).

**Figure 6:**
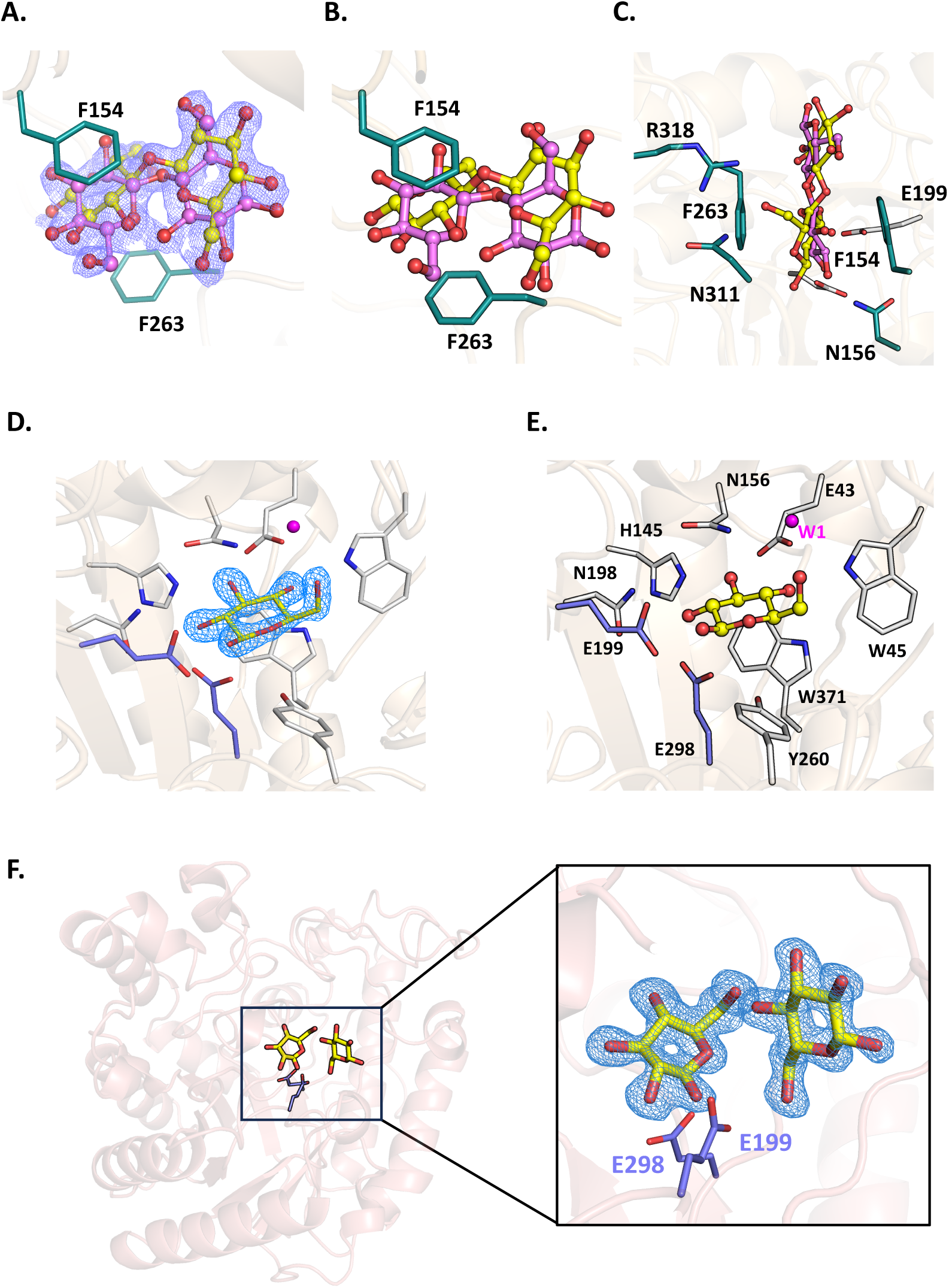
Interactions of AoBgl with bound sugars. (A., B.) Cellobiose molecule shown as yellow and light magenta ball and sticks captured in a double conformation well-stacked between phenylalanine residues 154 and 263 shown as blue-green sticks. The 2*F_o_-F_c_* electron density map contoured at 1σ level is shown around the cellobiose molecule. (C.) Bound cellobiose at the entry of the active site crater is also stabilized by polar interactions as well as stacking interactions with the residues lining the crater. (D., E.) Glucose at the active site stabilized by polar interactions with residues shown as light grey sticks and the catalytic glutamates shown as blue sticks. There is also a water molecule (W1) interacting with the glucose. The 2*F_o_-F_c_* electron density map contoured at 1σ level is shown around the glucose molecule. (F.) Overall view of the glucose-bound AoBgl structure. Two glucose molecules have been captured at its active site. Inset: zoomed-in view of the active site with catalytic glutamates shown as blue sticks and the glucose molecules are shown as yellow sticks. The 2*F_o_-F_c_* electron density map contoured at 1σ is shown as a sky-blue mesh around the glucose molecules.

Interestingly, a molecule of β-D-glucose detected at the active site formed strong hydrogen bonding interactions with the catalytic glutamate E298. The glucose molecule might have been present as a contaminant in the cellobiose solution which was used for soaking the crystal of apo-AoBgl. The bound glucose at the active site displaces the water molecules, which form hydrogen bonds with the nucleophilic glutamate residue. Glucose at the active site is further stabilized via hydrogen bonding interactions with N198, H145, N156, E43, W45, W371 and Y260. It also forms polar interactions with the water molecule indicated as W1(Figures 6D, E). The water interacting with glucose has a temperature B factor of 11.96 Å^2^, indicating that the water is well ordered at that site, and it might be essential in stabilizing the bound molecule of glucose. The two sugar moieties in cellobiose, together with one molecule of glucose, bound at the active site, the three glucose binding sites at the catalytic crater of AoBgl. A second molecule of glucose has been observed to be bound towards the entrance of the catalytic crater (Figure S6, S7). The binding of this glucose molecule at this position might indicate that AoBgl has alternate sites for glucose binding that might aid in keeping the environment around the active site residues glucose-free; thereby, such a site would contribute towards its high glucose tolerance. The structural details of AoBgl in both apo and sugar-bound state are described in supplementary movie M1 and Figure S6 (see supporting information).

We also observed a change in the conformation of a loop region ranging from amino acid residues 323-327 (Figure 7A, B). The major change is observed for the positions of residues A325 and D326. The conformational change in this region also involves a tyrosine residue at the 323^rd^ position that lines the opening of the active site crater. This tyrosine (Y323) is a part of the +2 site which plays a key role in sugar binding in glycosyl hydrolases. The loop region forms a flap-like structure, which is in “open” form in the apo structure. However, it “closes” on oligosaccharide binding. The net displacement, which is determined to be around 3.2Å, can be one of the ways by which the enzyme regulates itself on sugar binding, and it might be an important determinant of the mode of sugar binding.

**Figure 7:**
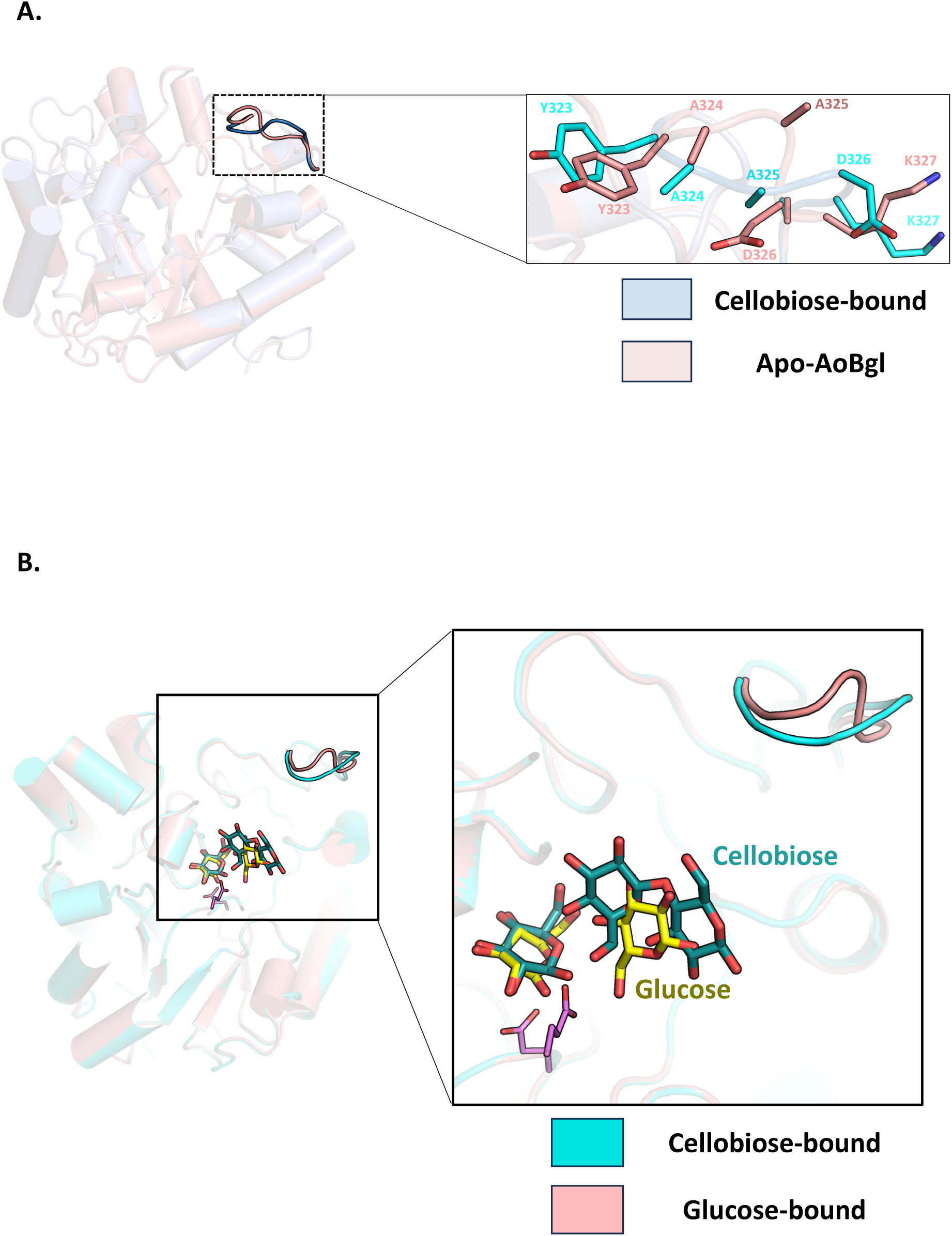
Dynamics observed in the loop region ranging from residues Y323-K327 due to cellobiose binding. (A) Superposition of apo-AoBgl (salmon) with the cellobiose-bound structure (light blue) shows change in conformation of residues in the loop (Y323-K327). (B) Superposition of glucose-bound (salmon) and cellobiose-bound (cyan) structures of AoBgl shows identical change in the loop conformation. Thereby indicating that the loop senses oligosaccharide ligands inside the catalytic crater.

In order to investigate whether the dynamics in the loop region (Y323-K327) has any role in sensing the type of sugar binding AoBgl, we solved the crystal structure of AoBgl soaked with 1.4 M glucose. The structure was solved at a high resolution of 1.2 Å. Two molecules of glucose were captured inside the active site crater (Figure 6F). These glucose molecules were found to occupy the same sites as seen occupied by sugar moieties in the cellobiose-bound structure thereby indicating a conservation in the sites for sugar binding. Interestingly, the loop region that has been discussed above has identical conformation in the glucose-bound form of AoBgl as seen in the apo-form. The conformational change is only seen in the loop when the disaccharide cellobiose binds in the catalytic crater while there is no such change on glucose binding alone. This is a strong indication of the role of this loop in sensing the type of sugar approaching its active site. As the monosaccharide, glucose cannot be further hydrolysed by the enzyme, the loop behaves in the same manner as the apo form. However, oligosaccharide binding triggers the loop closure which sends a message regarding the substrate approach towards the catalytic crater of AoBgl.

### Selectivity of AoBgl towards different oligosaccharides

Molecular Dynamics (MD) simulations were performed to understand the selectivity of substrates that AoBgl can hydrolyse as GH5 enzymes are known to be active on a variety of different sugars. The crystal structure for the apo form was used to carry out simulations of the apo form, whereas the cellobiose complexed structure was used as a starting structure for carrying out MD simulations with laminaribiose and laminaritriose (Figure 8A.) bound forms of the enzyme. All the simulation runs were initially carried out for 100 ns. However, the simulation of the laminaritriose-bound form was extended for another 400 ns (500 ns run in total) to check for stability and residence period of sugar at the active site of AoBgl.

**Figure 8:**
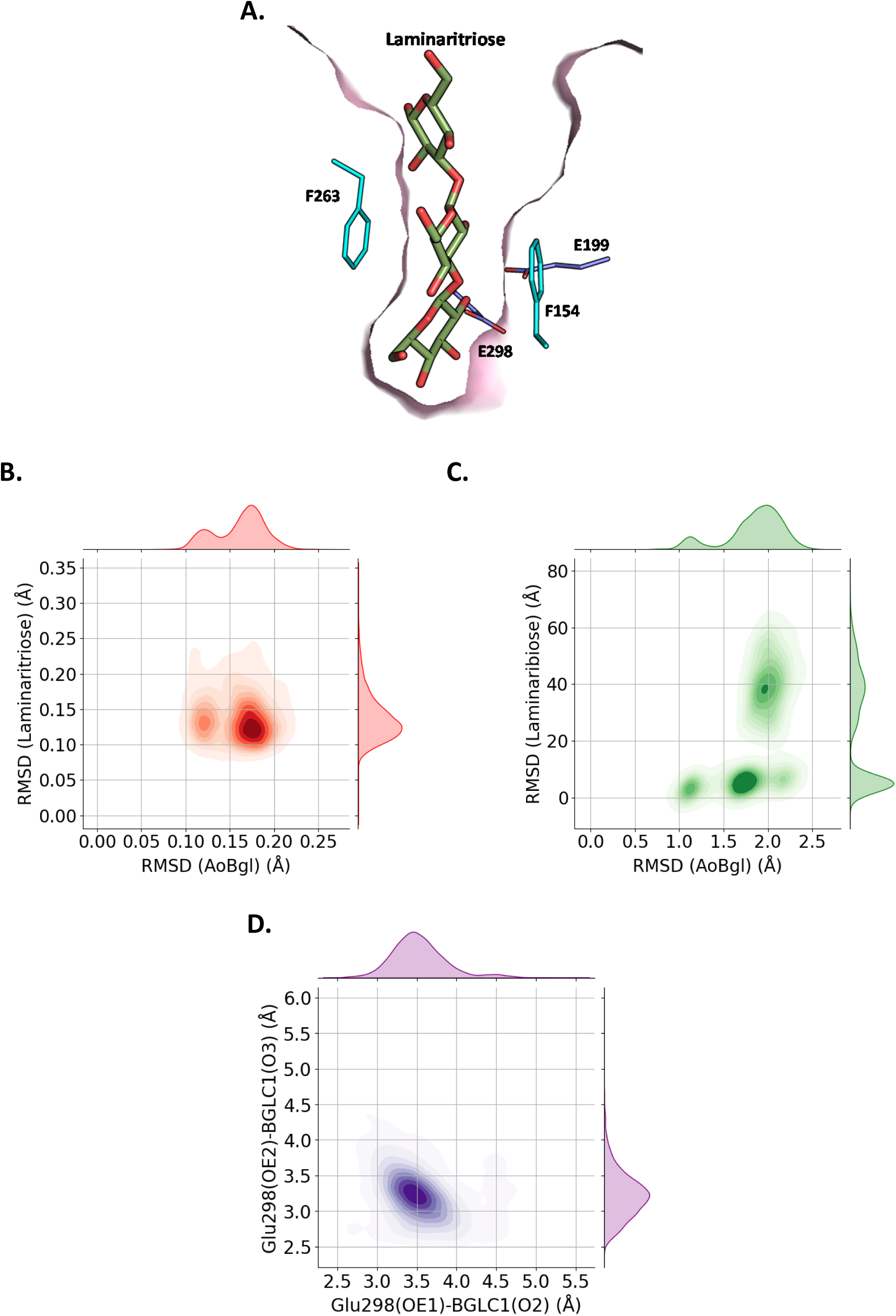
Insights into the sugar recognition by AoBgl using MD simulations. (A.) Binding pose of laminaritriose inside the catalytic crater of AoBgl The phenylalanine residues F154 and F263 which form stacking interactions to stabilize the bound ligand are shown as cyan sticks and the catalytic glutamates are shown in dark blue sticks. (B.) 2D plot of RMSD of laminaritriose vs RMSD of AoBgl during MD simulations. Laminaritriose shows only one population with RMSD of around 0.15 Å, indicating it is very well stabilized inside the catalytic crater of AoBgl. (C.) 2D plot of RMSD of laminaribiose vs RMSD of AoBgl during 100 ns of MD simulation run. Multiple populations of laminaribiose can be observed, which are around 40Å, which indicates that it is not stable and moves away from the protein’s catalytic pocket. Populations of AoBgl are within 1-1.5 Å which shows that the protein is stable throughout the duration of the simulation run. (D.) 2D plot of distance between OE2 of E298 in AoBgl and BGLC1(O3) in the bound laminaritriose vs OE1 of E298 in AoBgl and BGLC1(O2) in the bound laminaritriose shows clearly that there is one major population of variables within hydrogen bonding distances that indicates there is a constant interaction between the ligand and the catalytic nucleophile throughout the duration of MD simulation run.

RMSD and the Root Mean Square Fluctuation (RMSF) were calculated to understand the overall stability of both apo and the complex forms of AoBgl. The RMSD curve indicates that the system is stable throughout the 100 ns of the simulation run (Figure S8). Distance between the CD of the two catalytic glutamates (E199 and E298) remains around 5.5Å throughout 100 ns of the MD simulation run (Figure S9) which indicates that AoBgl is expected to show the retention mechanism of sugar hydrolysis.

The RMSF curves, however, were very distinct for each of the three substrates that were complexed with AoBgl. There were very high fluctuations in several regions in the disaccharide-complexed structures compared to the apo form indicating that the system is destabilized by the binding of this ligand (Figure S10). However, for the laminaritriose bound form, it was observed that the fluctuations were significantly decreased and the RMSF curve resembled that of the apo structure. This indicates that laminaritriose can be a substrate for hydrolysis by AoBgl, whereas laminaribiose is not. The RMSF curves show that the catalytic pocket of AoBgl can accommodate trisaccharides more effectively than disaccharides. Disaccharide molecules like laminaribiose were found to escape the pocket of AoBgl within 20-40 ns of MD simulation run, indicating that it cannot bind to the catalytic pocket effectively. This can also be seen from the crystal structure of the cellobiose-soaked form of AoBgl which indicates that three sugar units can be accommodated inside the catalytic pocket of the enzyme.

To further confirm our claim that laminaritriose interacts stably with the catalytic residues of AoBgl, the simulation run was extended by another 400 ns to check if the interactions were disrupted. The 2-dimensional plots in Figures 8 B. and C. compare the RMSD of the laminaribiose and laminaritriose ligands with that of AoBgl, respectively. It is clear from these plots that laminaritriose is stabilized better inside the crater of AoBgl whereas laminaribiose escapes from the active site, which is indicated by the high values of RMSD for the ligand (supplementary movies M2 and M3). Maintenance of continuous polar interactions between BGLC1-O2 of laminaritriose with OE1 of the catalytic nucleophile (E298) and BGLC1-O3 of laminaritriose with OE2 of E298 was observed throughout the duration of the MD simulation run. The 2-dimensional plot compares the distances between the oxygen atoms in glucose units in laminaritriose and oxygen atoms in the catalytic nucleophile E298 throughout the duration of MD simulation run (Figure 8D). The importance of the catalytic nucleophile E298 has been further confirmed by the mutation of the glutamate residue to serine (AoBgl-E298S). The AoBgl-E298S mutant has been expressed and purified. Biochemical assays performed with this purified AoBgl-E298S catalytic mutant of AoBgl do not indicate hydrolysis on any of the substrates (p-NPG, laminaritriose, or laminarin) which confirms our claim from the MD-simulation studies that the interaction between laminaritriose and the catalytic nucleophile is important for the enzyme to be functional towards the bound substrate.

The structural superposition of our cellobiose-bound AoBgl structure with the exo-β-(1,3) glucanase from *C. albicans* with bound laminaritriose (PDB ID:3N9K) (39), shows that the sugar binding sites are conserved between the two proteins. The crystal structure with bound cellobiose and glucose can represent the actual mode of sugar binding inside the catalytic crater of AoBgl. We have observed that in the cellobiose complexed structure of AoBgl, the bound cellobiose molecule inside the catalytic crater is very well stacked between F154 and F263 residues which are also the gatekeeper residues of the catalytic crater (Figure 8A.). The mutation F263A shows total loss of laminarin hydrolysis activity of the enzyme which further supports our claim that the stacking interactions of the sugar with the aromatic amino acids like phenylalanine inside the active site crater is essential for the catalytic activity of AoBgl.

## Discussion

Utilization of algal biomass to generate bioethanol is a promising and relatively hassle-free source of green energy. This process, however, requires a separate set of enzymes that hydrolyse β-(1,3) linkages of sugars predominant in algal cell walls. The structural and biochemical characterizations of such enzymes are relatively few in existing literature thus, more research is needed towards their industrial applications.

In this study, we have characterized one such recombinant exo-β-(1,3) glucanase from *A. oryzae* (AoBgl). Purification of the recombinantly expressed AoBgl required three steps of chromatographic techniques as it exhibited poor binding with the Ni-NTA column. Anion exchange chromatography at pH 5.2 could improve the purity significantly owing to the low pI of AoBgl. In the next step, size exclusion chromatography could remove all the remaining minor impurities and AoBgl was eluted under a single peak as a monomeric protein. This agrees with previous studies on exo-β-(1,3) glucanases which are also monomers (37, 40, 41). Our study reports the first successful recombinant expression and purification of AoBgl from a bacterial expression system.

At the initial stages of this study, we performed enzymatic assays of AoBgl using substrates for β-glucosidases (both p-NPG and cellobiose) as a previous report on this enzyme purified from native source stated its ability to hydrolyse disaccharides with β-(1,4) linkages (29). To our surprise, purified recombinant AoBgl hydrolysed only p-NPG but, there was no detectable cellobiose hydrolysing activity for this enzyme. We would like to emphasise that this as an important observation as several published literature characterized β-glucosidases based on p-NPG hydrolysis activity only (42–45). Our study shows that p-NPG hydrolysis cannot be considered to mimic enzymatic hydrolysis activity on real disaccharide substrates like cellobiose.

Several glycosyl hydrolases with conserved overall structural fold (30, 34, 41, 42, 46, 47) might hydrolyse p-NPG; however, hydrolysis of the actual substrate might require proper positioning of the catalytic motifs for the activity. The inability hydrolyse cellobiose confirms that AoBgl is not a β-glucosidase that can act on β-(1,4) linkages. This prompted us to investigate sequences and possible structures of homologs of AoBgl which confirmed that it belongs to GH5 family, and its actual substrates are β-(1,3) linked sugars. The hydrolysis assays with p-NPG revealed that 55°C was the optimum temperature for p-NPG hydrolysis. A previous study has reported that AoBgl recombinantly expressed in fungal expression systems do not show activity beyond 50°C (48). However, when expressed in bacterial systems, the enzyme shows good activity as well as stability at 50°C. AoBgl has been found to hydrolyse laminarin which is a polysaccharide with β-(1,3) linked glucose chains effectively. This makes it a promising candidate for further exploration concerning its use in commercial bio-ethanol production process from algal biomass. The kinetic characterization showed a deviation from Michelian behaviour for laminarin hydrolysis; this might be because laminarin, being a branched-chain polysaccharide, possesses multiple ends for the AoBgl to act on. This might lead to more than one molecule of the enzyme hydrolysing one molecule of laminarin simultaneously which would cause deviation from standard Michelian kinetic behaviour. The carbohydrate selectivity of AoBgl has been further confirmed by performing hydrolysis assays with laminaritriose. The fact that AoBgl successfully carried out laminaritriose hydrolysis indicates its preference towards β-(1,3) linked oligosaccharides.

To further investigate the molecular determinants differentiating between the enzymes that can hydrolyse β-(1,3) and β-(1,4) linkages, we have solved, for the first time, the high-resolution crystal structures of AoBgl. The TIM barrel fold observed for AoBgl is also the structural feature present in several other glycosyl hydrolases, especially those belonging to families GH1 and GH5 (34, 37, 39). Although the structural fold of AoBgl was identical to both GH1 and GH5 family of enzymes, structure-based comparisons revealed that it shared much higher levels of similarity with GH5 enzymes than their GH1 counterparts. These structural as well as sequence-based comparisons made it clear that it aligned better with GH5 family than members of GH1 family of enzymes (Figure 2, S2, S3 and S4).

To understand the mode of sugar binding at the active site crater of AoBgl, one crystal of apo-AoBgl was soaked in a solution of cellobiose. As mentioned in the previous sections, AoBgl did not show cellobiose hydrolysis, this made it a good disaccharide to study the interactions at the active site crater as it would bind to the enzyme but would not be hydrolysed by it. A previous study has shown how a bound trisaccharide ligand interacts with a homologous exo-β-(1,3)-glucanase (39), but to capture the actual substrate, an inactive mutant enzyme was used for crystallization. We wanted to understand how the residues at the catalytic crater arrange themselves to stabilize a sugar bound to the wild-type enzyme itself. We could capture cellobiose at the entrance of the active site crater well-stacked between the gate-keeper residues, F154 and F263. The cellobiose solution used for soaking the AoBgl crystal had around 33mM glucose as a contaminant; because of which we could observe the electron density corresponding to a glucose molecule inside the active site crater of the enzyme. The two glucose moieties in cellobiose along with one bound glucose molecule at the catalytic crater prove that AoBgl can accommodate three sugar units at its active site crater. Structural alignment with the mutant exo-β-(1,3)-glucanase from *C. albicans* with bound laminaritriose (PDB ID: 3N9K) (39) shows that the sites inside the active site crater which interacts with the bound ligand are identical in both the cases. Therefore, the cellobiose and glucose-bound AoBgl structure might depict the actual mode of substrate binding in its catalytic crater.

We have also observed that a loop region comprising residues Y323-K327 shows a change in conformation on ligand binding, which has not been seen in any previously reported structures of enzymes that share sequence identity with AoBgl (37, 39, 47). This loop region forms a part of the entrance to the catalytic crater and such changes in conformation might relate to sensing of the disaccharide ligand binding to the enzyme. The role of this loop in sensing the type of sugar has been further validated structurally from the high-resolution glucose-bound structure of AoBgl. The loop maintains a conformation that is identical to the apo-AoBgl in the glucose-bound form. Glucose-bound AoBgl does not exhibit a “closing” of the loop as was observed in the cellobiose-bound form of the enzyme. This clearly shows that the loop can sense the type of sugar moiety approaching the catalytic crater of the enzyme. One previous structural study using an inactive mutant of homologous enzyme of AoBgl does not reflect such changes in the said loop on ligand binding. This might also indicate that the interactions with the bound ligand in the wild-type enzyme can differ from those in the mutant. Our study is the first attempt aimed at investigating the interactions of a bound disaccharide ligand at the active site crater of a wild-type enzyme of related functionality. Insights obtained from correlating the apo, cellobiose-bound, and glucose-bound structures indicate that the loop region undergoes conformational changes depending on the type of sugar moiety approaching the enzyme, thereby having a role in sensing its chemical and oligomeric nature.

The insights obtained from molecular dynamics (MD) simulations provided an idea about the oligomeric nature of sugar molecules that can bind, interact stably, and ultimately serve as a suitable candidate for hydrolysis by AoBgl. The information regarding the binding site of sugar inside the catalytic crater of the enzyme was obtained from the cellobiose-bound structure. As mentioned in the previous sections, there was a molecule of glucose captured along with cellobiose at the catalytic crater thereby making the catalytic crater accommodate three sugar moieties (Figure S8). When compared with the exo-β-(1,3) glucanase from *C. albicans* with bound laminaritriose (PDB ID: 3N9K), we could observe that the same sites were occupied with the sugar moieties of laminaritriose thereby leading to the conclusion that the sugar binding sites of AoBgl as obtained from our crystal structure is identical to what has been previously reported in enzymes of this family. The laminaritriose molecule bound structure of AoBgl was prepared and simulated for 500 ns to check for the residence period of the bound sugar. It was observed that laminaritriose maintained stable hydrogen bonding interactions with the catalytic glutamate E298 throughout 500 ns of the simulation run. The trajectories analysed from the laminaribiose bound structure revealed that the sugar escapes from the catalytic crater within 30-40 ns which means that it is not stabilized within the crater, and this is also supported by the RMSF plots that indicate greater degrees of fluctuations within the system in laminaribiose bound form whereas the system is much more stable in the laminaritriose bound form. This evidence indicates that laminaritriose, a trisaccharide is the shortest sugar unit that AoBgl might hydrolyse. This is further corroborated by laminaritriose hydrolysis assay which shows that AoBgl can bring about laminaritriose hydrolysis. Analysis of the trajectories generated after 500 ns MD simulation run for the laminaritriose-bound form of AoBgl revealed that the bound ligand is well-stacked between the F154 and F263 residues throughout the run. To experimentally validate this observation, mutation of the phenylalanine residue at 263^rd^ position to alanine was carried out and the resultant mutant AoBgl-F263A was expressed, and it was found that it was inactive against laminarin. This proves that the binding of laminaritriose, a trisaccharide at the catalytic crater is the same as that of the polysaccharide laminarin.

## Conclusion

The detailed biochemical and structural investigations carried out in the above study provides molecular insights into the functioning of a fungal exo-β-(1,3) glucanase (AoBgl). AoBgl was found to carry out hydrolysis of the polysaccharide laminarin to produce glucose. In addition to that, the enzyme very low levels of product inhibition. These properties make it a very promising candidate for application in the industrial sector for production of third generation bio-ethanol from algal biomass. We also solved the crystal structure of AoBgl at near-atomic resolutions which aided us in understanding the mode and sites of sugar binding inside the catalytic crater of the enzyme. These high-resolution structures have also helped us to identify a loop region which shows conformational changes only when oligosaccharides bind to the crater. This is a novel finding as previously characterized proteins with identical structural fold have not shown such dynamics on oligosaccharide binding. These structural insights have helped us to infer that the loop residues might have a role in sensing the approach of substrate inside the crater of the enzyme. In addition to these, MD simulation-based studies have shown that trisaccharides are better stabilized in the catalytic crater of AoBgl than disaccharides. This indicates that trisaccharides like laminaritriose can be the shortest sugar moiety that can be accommodated stably and subsequently hydrolysed by AoBgl. Our characterizations of this enzyme would prove helpful in structure-guided protein engineering experiments in future for further improvements in its catalytic performance to be considered an ideal candidate for industrial saccharification processes.

## Experimental procedures

### Expression of recombinant AoBgl

Gene (*aobgl)* coding for AoBgl was cloned in pET43a vector between *NdeI* and *NcoI* restriction sites. The construct was designed in such a way that the expressed polypeptide of AoBgl contains a hexahistidine tag at the C-terminus. The recombinant expression plasmid was transformed into *Escherichia coli* BL21(DE3) cells. The cells were grown on Luria-Bertani (LB) media at 37°C until the OD_600_ reached 0.8. The expression of AoBgl was induced by addition of 0.1mM IPTG and protein expression was done for 16 hours at 22°C.

After the said incubation time, the cells were harvested by centrifuging at 8000 r.p.m for 8 minutes at room temperature. The cells pellets were resuspended in lysis buffer (50 mM Tris-Cl, 400 mM NaCl, pH 7.4) and the cells were lysed by sonication. The sonication conditions used were 3 seconds on and 1 second off at 40% amplitude. Following lysis, the soluble and insoluble fractions were separated by centrifugation at 16000×g at 4°C for 30 minutes. The protein concentration of the supernatant was measured by the Bradford method of protein estimation (49). The presence of soluble AoBgl in the crude extract was confirmed by protein content analysis on a 12% SDS-PAGE.

### Purification of AoBgl using chromatographic techniques

The culture for AoBgl expression was scaled up to 6 litres and the conditions mentioned above was used to treat the cells to obtain the protein in soluble fractions. The cell free extract that was obtained post lysis was purified by Ni-NTA affinity chromatography. Fractions obtained after Ni-NTA purification were not pure enough as AoBgl co-eluted with several contaminants. The fractions obtained after Ni-NTA purification were pooled and diluted 10-fold to reduce salt concentration. The buffer used for dilution was 50 mM sodium acetate, pH 5.2. The dilution caused several contaminant proteins to precipitate, and the solution obtained after filtration was loaded on DEAE sephadex anion exchange column for further purification. The fractions eluted after anion exchange chromatography had a much higher degree of purity. These were further pooled, concentrated, and purified using size exclusion chromatography with a Superdex-75 column. Extremely pure AoBgl was obtained, which was free of most contaminants. This highly pure form of the protein was used for performing biochemical characterizations and crystallization. All the chromatographic purification steps were carried out using ÄKTA Pure FPLC system (GE Healthcare).

### Biochemical characterizations of AoBgl

AoBgl purified after gel filtration chromatography was characterized biochemically. Assays were performed using p-NPG, which is the artificial substrate for enzymes that hydrolyse glycosidic bonds. It has been widely used for biochemical characterization and activity measurements of these classes of enzymes. Assays were carried out by incubating 50 µl of 40 mM p-NPG in 440µl 50mM sodium acetate buffer of pH 5.5 and 10 µl of AoBgl was added to the reaction mixture. It was incubated for 10 minutes and after that 500 µl of Na_2_CO_3_ was added to it. The yellow colour obtained was quantified spectrophotometrically by measuring absorbance at 405 nm. All biochemical assays on AoBgl using p-NPG as substrate has been performed in the same manner.

After the determination of optimum temperature and pH for enzyme activity using p-NPG, laminarin, the actual substrate for AoBgl was also used for biochemical assays of the enzyme. Suitable concentration of the purified enzyme was incubated with laminarin in 50 mM sodium acetate buffer of pH 5.5 at 50°C for 30 minutes, after which the reaction was stopped by incubating the mixture at 95°C for 10 minutes. After enzyme inactivation, the amount of glucose produced by laminarin hydrolysis was quantified spectrophotometrically at by measuring the absorbance at 505 nm using a GOD-POD kit (Accurex). Hydrolysis assays of AoBgl with laminaritriose, cellotriose and cellobiose have been performed in the same manner.

#### Determination of temperature optima

To determine the optimum temperature for hydrolysis by AoBgl, the assays were carried out in 50 mM sodium acetate buffer of pH 5.5. The reaction mixtures were prepared as described above. The reaction mixtures were incubated at temperatures of 25°C, 30°C, 35°C, 40°C, 45°C, 50°C, 55°C, 60°C and 70°C, for 15 minutes for equilibration. After which 10µl AoBgl was added, and the assays were carried out as described in the previous section.

#### Determination of pH optima

Optimum pH of hydrolysis by AoBgl was assayed in buffers of various pH ranging from 3 to 8. 50 mM sodium acetate buffer was used for assays in pH range 3 to 5.5, 50 mM sodium phosphate buffer was used for assays in pH range of 6 to 7.5 while 50 mM Tris-Cl buffer was used for assays at pH 8. 40 mM p-NPG, which was used as substrate, was also dissolved in the same buffers. The optimum pH for laminarin hydrolysis was also determined by carrying out assays with laminarin as substrate at pH range 3.0-8.0.

The reaction mixture comprised of 50µl of 40 mM p-NPG in 440µl of buffers of the pH range 3 to 8. The reaction mixture was incubated at 55°C for a period of 15 minutes. After the lapse of the said time, 10 µl AoBgl was added, and the assays were carried out. in the same way, as mentioned in the previous section.

#### Determination of product inhibition

To determine the inhibitory effects of the product, glucose, the enzyme assays of p-NPG hydrolysis were carried out by incubating the enzymes at increasing glucose concentrations. A 2 M stock solution of glucose was prepared in a 50 mM sodium acetate buffer of pH 5.5. 50µl of 40 mM p-NPG was incubated with varying glucose concentrations ranging from 0-2M. Each reaction mixture was incubated for 15 minutes at 55°C. After that, 10µl of the enzyme was added and the mixtures were incubated for 10 minutes. Next, the reaction was stopped by adding 500µl of 0.2M sodium carbonate and the yellow colour generated was assayed spectrophotometrically by measuring the absorbance at 405 nm.

#### Enzyme kinetics studies

Suitable dilutions of the enzyme were prepared with the reaction buffer (50 mM sodium acetate buffer of pH 5.5) for performing the assays involving hydrolysis of p-NPG. Kinetics studies were performed to determine the K_M_ and V_Max._ For AoBgl, p-NPG was used as substrate. The reaction mixtures were prepared with increasing concentrations of p-NPG (0.1-40 mM) and incubated at 55°C for 15 minutes. After which 10µl of suitable dilution of AoBgl was added and the reaction was allowed to continue for 10 minutes, after which the assays were performed as mentioned in the previous section. Enzyme kinetics for AoBgl were also performed using the actual substrate, laminarin. AoBgl was incubated with varying concentrations of laminarin (0.05-47.5 mg/ml) at 50°C for 30 mins. The determination of enzyme activity was carried out in the same manner as described in the previous section.

#### Assessment of stability of the enzyme at different temperatures

AoBgl was initially incubated at a temperature of 55°C; however, the enzyme activity was lost after 1 hour of incubation. Following this, the incubation was done at 30°C. To assay the relative activity, aliquots of 10 µl of the enzyme were made after every 12 hours, and the activity assays were done in the same manner as described in section 2.4. Assays were done up to 48 hours of incubation and activities relative to that of the zero-hour reading were plotted using Graph Pad Prism 8.4.2. Tryptophan fluorescence was also recorded by incubating AoBgl at temperatures from 30-60°C. The experiment was carried out to determine the thermal unfolding profile of AoBgl. Details of this experiment are mentioned in Supporting Information.

### Crystal structure determination of AoBgl

#### Crystallization of AoBgl

The purified AoBgl was concentrated up to 13 mg/ml for setting up the crystallization screens. The crystallization drops were set up using commercially available screens such as JCSG+ (Qiagen), PEG suite (Qiagen), PEG-Rx (Qiagen), Index (Molecular Dimensions), and PEG Ion (Qiagen) by mixing 2μl of AoBgl with 2 μl mother liquor equilibrated against 50 μl of the same reservoir solution. The screens were set up using a Phoenix (Art Robins) crystallization robot at the Protein Crystallography Facility, Indian Institute of Technology Bombay, India. The sitting drop vapor diffusion method was used for crystallization of AoBgl and the crystallization trays were incubated at 18°C for six months for crystals to appear. Crystals of apo-AoBgl were obtained in mother liquor having 1.6 M sodium citrate tribasic dihydrate (pH 6.5) and 0.2 M sodium chloride with 20% PEG.

#### X-ray diffraction data collection and processing

X-ray diffraction of the AoBgl crystals was performed to elucidate the structure of the enzyme. Diffraction experiments were performed under liquid nitrogen cryoconditions (100K). The mother liquor that formed the crystals was 1.6 M sodium citrate tribasic dihydrate (pH 6.5). The mother liquor itself was a cryoprotectant. A crystal of AoBgl was picked from the mother liquor using a nylon loop, and flash frozen in the liquid nitrogen stream at 100 K. A diffraction data set was collected by the rotation method with 0.5° rotation per frame at a wavelength of 1.5418 Å using Cu *K*α X-ray radiation generated by a Rigaku Micromax 007HF generator having R-Axis IV++ detector at the Protein Crystallography Facility, IIT Bombay.

AoBgl crystals were soaked with cellobiose to obtain structural information regarding the mode of binding of disaccharide sugars to the active site crater. A solution of 200 mM cellobiose was prepared in the cryoprotectant (which is the mother liquor) and one AoBgl crystal was soaked in that solution for 10 minutes. Then the crystal was picked up using a nylon loop, and flash-frozen in the liquid nitrogen stream at 100 K. A diffraction dataset was collected from the cellobiose-soaked crystal using the same procedure as described above using home X-ray source.

For obtaining insights into the mode of binding of monosaccharides in the catalytic crater glucose complexed AoBgl structure was determined. One AoBgl crystal was soaked with 1.4M glucose solution having 30% glycerol as the cryo-protectant. Then, it was immediately fished out using a nylon loop and flash-frozen in liquid nitrogen. The frozen crystal was shipped to the synchrotron radiation facility. The diffraction experiment and data collection for the glucose-soaked crystal was carried out by rotation method with oscillations of 0.5° per frame at PX-BL21 beamline in the Indus-2 synchrotron at Raja Ramanna Centre for Advanced Technology (RRCAT), Indore, India.

Indexing, integration, and scaling of all the datasets were performed using XDS (50). F2MTZ and CAD modules of CCP4 (51) were used for converting the intensities to structure factor. Table 1 presents the data collection statistics.

#### Structure determination, model building, and refinement

The structure of apo-AoBgl was solved using molecular replacement (MR) method and the structure (PDB ID: 1CZ1) of an exo-β-(1,3) glucanase from *Candida albicans* (37) was used as a template. This template structure was selected as it had the highest sequence identity of 46.9% with the AoBgl sequence and the query coverage was also satisfactory. MR performed using PHASER (52) program of CCP4 (53) could find one molecule of AoBgl in the asymmetric unit. After the phasing, the model was built using BUCCANEER (53) program of CCP4 suite and manually by visual inspection using COOT (54). The refinement of the complete model of apo-AoBgl was carried out using REFMAC5 (55).

The structures of the AoBgl-cellobiose complex and AoBgl-glucose complex were solved separately by molecular replacement using the coordinates of apo enzyme. First few cycles of refinement of only the protein molecule were performed by REFMAC5. Analysis of the *F*_o_−*F*_c_ electron density map revealed presence of cellobiose and glucose molecules in the respective complexed structures. Subsequently, the sugar molecules were placed inside the *F*_o_−*F*_c_ electron density map and the structure was refined further. The refinement statistics for all the structures are presented in Table 1.

### MD simulation studies

To determine the types of molecules that are hydrolysed by AoBgl, and to understand the mode of binding and dynamics, molecular dynamics (MD) simulation studies were performed. The apo and cellobiose-bound structures were selected for these studies. The apo structure was prepared using the solution builder module in CHARMM-GUI (56). For substrate-bound forms, the cellobiose-bound structure was used. Laminaribiose, laminaritriose molecules were added by structural alignment of AoBgl with an exo-β-(1,3) glucanase from *C. albicans* in complex (PDB ID: 3N9K) with laminaritriose (PDB ID: 3N9K) (39).

Molecular dynamics simulations of all the structures were performed using Gromacs 2022.2 with CHARMM36 (57) force field. The macromolecular simulation system was prepared with help of CHARMM-GUI (56) server during which protein-ligand complexes were placed in a cubic box of water molecules, ensuring a minimum distance of at least 10 Å between the protein and the box edges. Appropriate counterions (such as Na+ or Cl-) were added to achieve a desired physiological ionic strength of 0.15 mol/L. The system was then energy-minimized to remove any steric clashes or close contacts. The system was subjected to several stages of equilibration, including NVT (constant number of particles, volume, and temperature) and NPT (constant number of particles, pressure, and temperature) ensembles. During equilibration, the temperature was controlled using the Nose-Hoover thermostat (58, 59), and the pressure was controlled using the Parrinello-Rahman barostat (60). Non-bonded interactions were calculated and the particle–mesh Ewald method was used to compute the long-range electrostatic interactions by specifying proper PME grid size. The SHAKE (61) algorithm was used to constrain all bond lengths involving hydrogen atoms which permits a 2 fs time step and the value of steps per cycle (time steps per cycle) was 10. After equilibration, the production MD simulations were carried out for 100 ns simulation time where atomic coordinates were captured every 10 ps. Properties of interest such as root-mean-square deviation (RMSD), root-mean-square fluctuation (RMSF), hydrogen bond interaction were calculated on the simulation trajectories. Visualization of trajectories were carried out using VMD and graphs were plotted using Graph Pad Prism 8.4.2.

### Site-directed mutagenesis of AoBgl

Site-directed mutagenesis was performed on AoBgl to confirm the role of residues towards stabilization of the bound sugar substrates catalyzing and their hydrolysis. The phenylalanine residue at the 263^rd^ position, which was found to form stacking interactions with the bound sugar substrate in the crystal structure, was mutated to alanine (AoBgl-F263A). Further, the glutamate residue at the 298th position, which was identified as the catalytic nucleophile from sequence and structure-based comparisons, was mutated to serine (AoBgl-E298S) in order to verify its role as the catalytic residue. The mutations were carried out using inverse-PCR with the mutation incorporated in the semi-overlapping primers itself. Primer sequences are given in Table S1. After the whole plasmid was amplified by PCR, the wild-type template was digested using DpnI, followed by transformation into *E. coli* DH5α cells. Each of the mutations was confirmed using sequencing. The mutants were expressed and purified in the same manner as the wild-type enzyme. Hydrolysis assays were performed using p-NPG, cellobiose, laminaribiose and laminarin using the same protocol as described in the sections above.

## Data availability

Most of the data are contained within the article and supporting information section. The atomic coordinates, as well as the structure factors for AoBgl-Apo (PDB ID: 8Z2W), AoBgl-cellobiose-bound (PDB ID: 8Z2X) and AoBgl-glucose-bound (PDB ID: 8Z2Y) structures, were deposited in the Protein Data Bank (http://wwpdb.org/).

## Supporting Information

This article contains supporting information.

## List of abbreviations

p-NPG: paranitrophenyl-β-D-glucopyranose
IPTG: isopropyl-β-D-thiogalactopyranoside
Ni-NTA: nickel-nitrilotriacetate
GH: glycosyl hydrolase

## Acknowledgements

We thank Dr. Ravindra Makde and Dr. Biplab Ghosh at the PX-BL21 beamline (BARC) at Indus-2, RRCAT, Indore, India for their support in diffraction data collection. We acknowledge the “Protein Crystallography Facility” at IIT Bombay. The work was supported by research funding to PB from the Department of Biotechnology, Government of India (grant number: BT/PR41982/PBD/26/822/2021).

