## Supporting Information for "Molecular insights into the oligosaccharide binding, recognition and hydrolysis by a fungal exo-β-(1,3)-glucanase"

**Short title :** Structural insights into sugar binding in a GH5 enzyme

*To whom correspondence should be addressed: Prasenjit Bhaumik, Department of Biosciences and Bioengineering, Indian Institute of Technology Bombay, Powai, Mumbai-400076, India; Telephone: +91-22-2576-7748; Fax: (+91)-22-2572-3480;

**Keywords:** GH5 enzyme, cellobiose, glucose, crystal structure, laminaribiose, laminaritriose, bioethanol, *Aspergillus oryzae,* laminarin, product inhibition

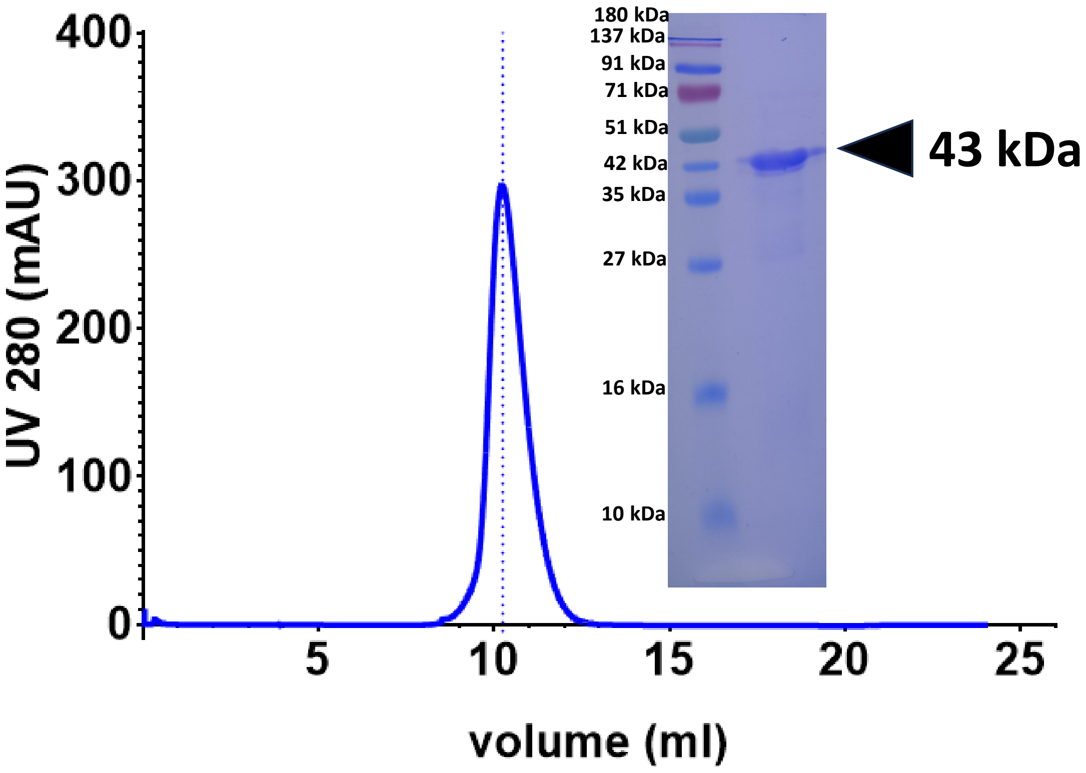


**Figure S1: Chromatogram after size exclusion chromatography of AoBgl showing single peak.** Inset: SDS PAGE showing single band corresponding to the molecular weight of AoBgl.

```
AoBgl    1  .............MLPLLLCIVPYCWSSRLDPRA..SSFDYNGEK.VRGVNLGGWLVLEP
NkBgl    1  .............MDVASSDTVYTFPPDEFKLGAATASYQIEGAWDENGKGPNIWDTLTH
HiBgl    1  MGSSHHHHHHSSGLVPRGSHMASMSLPPDFKWGFATAAYQIEGSVNEDGRGPSIWDTFCA
Td2F2    1  ....................MAGERFPADFVWGAATAAYQIEGAVREDGRGVSIWDTFSH
Bgl6     1  MG...........SMTSDTARSYRFPEGFLWGAATAAYQIEGSSMADGAGESIWDRFSH

AoBgl   45  WITPSIFD.AAGAEAVDEWSLTKILGKEEAEARLSAHWKSFVSAGDFQRMADAGLNHVRI
NkBgl   47  EHPDYVVDGATGDIADDSYHLY....................KEDVKILKELGAQVYRF
HiBgl   61  .IPGKIADGSSGAVACDSYKRT....................KEDIALLKELGANSYRF
Td2F2   41  .TPGKIADGTTGDVACDSYHRY....................GEDIGLLNALGMNAYRF
Bgl6    49  .TPGNMKDGDTGDVACDHYNRW....................REDIELMKRLNLQAYRF

AoBgl  104  PIGYWALGPLEG..DPYVDGQLEYLDKAVEWAGAAGLKVLIDLHGAPGSQNGFDNSGRRG
NkBgl   86  SISWARVLPEGH.DNIVNQDGIDYYNNLINELLANGIEPMVTMYHWDLPQALQDLG....
HiBgl   99  SISWSRIIPLGGRNDPINQKGIDHYVKFVDDLIEAGITPFITLFHWDLPDALDKRY....
Td2F2   79  SIAWPRIVPLGA..GPINQAGLDHYSRMVDALLGAGLQPFVTLYHWDLPQPLEDRL....
Bgl6    87  SVSWSRVIPQGR..GAINPKGLAFYDRLVDGLLEAGIEPLATLYHWDLPAALDDRG....

AoBgl  162  AIQWQQG.DTVEQTLDAFDLLAERYLGSDTVAAIEAINEPNI..................
NkBgl  141  ..GWP.N.LVLAKYSENYARVLFKNFG.DRVKLWLTFNEPLTFMD.GYASEIGMAPS...
HiBgl  155  ..GGFLNKEEFAADFENYARIMFKAI..PKCKHWITFNEPWCSAILGYNTG.YFAPGHTS
Td2F2  133  ..GW.GS.RATATVFAEYADIVVRQLG.DRVTHWATLNEPWCSAMLGYYLG.VHAPGHTD
Bgl6   141  ..GW.LN.PDIADWFADYGQVLFEKFK.GRVKTWGTINEPWVIVDGGYLHG.ALAPGHRS

AoBgl  203  ..............................................PGG...
NkBgl  192  ......INTPGIGDYLAAHTVIHAHARIYHLYDQEFRAEQGGKVGISLNINWCEPAT.NS
HiBgl  210  DRSKSPVGDSAREPWIVGHNILIAHARAVKAYREDFKPTQGGEIGITLNGDATLPWDPED
Td2F2  187  L.........KRGLEASHNLLLGHGLAVQAMRAAA..PQPLQIGIVLNLTPTYPAS.DS
Bgl6   195  A.........YEAVIAGHNVLRAHGAAVRRFREVG....EGQIGIVLNIEPKYPAS.DK

AoBgl  206  ...VD......QGKLQEYYGSVYGIVNKYNAGTSV..........VYGDGFLPVESWNGF
NkBgl  245  AEDRASCENYQQFNLGLYAHPIFTEEGDYPAVLKDRVSRNSADEGYTDS......RLPQF
HiBgl  270  PADIEACDRKIEFAISWFADPIYF..GKYPDSMRK..........QLGD......RLPEF
Td2F2  234  PEDVAAARRFDGFVNRWFLDPLAG..RGYPQDMLD..........YYGA......AAPQA
Bgl6   240  PEDEAARRRAEAQMNRWFLDPLMG..RGYPEELTD..........VYGA......AWREF

AoBgl  247  KTEGSKVVMDTHHYHMFDNGLIAMDIDSHIDA..........VCQFAHQHLEASDKPVIV
NkBgl  299  TAEEVEYIRGTHDFLGIN.FYTALLGKSGVEGYEPSRYRDS..GVI...LTQDAAWPISA
HiBgl  312  TPEEVALVKGSNDFYGMN.HYTANYIKHKTGV.PPEDDFLGNLETLFYNKYGDCIGPETQ
Td2F2  276  NPEDLTQIAAPLDWLGVN.YYERMRAVDAPDASLPQAQRLD...........DPDLPH.T
Bgl6   282  PKEDFELIAEPTDWMGLN.WYTRAVPENAPDAWPTRSRPVR...........QTQHAHTE

AoBgl  297  GEWTGA....VTDCAKYLNG...........KGNGARYDG..SYAADKAIGDCSSLATGFV
NkBgl  353  SSWLKVVPWGFRKELNWIKNEY.NNPPVFITENGFSDYG.......GLND......TGR
HiBgl  370  SFWLRPHAQGFRDLLNWLSKRY.GYPKIYVTENGTSLKGENDMPLEQVLED......DFR
Td2F2  323  ADR.EVYPEGLYDILLRLHNDY.PFRPLYITENGCALHD..EIAEDGGIHD......GQR
Bgl6   330  TGW.EVYPPALTDTLVWLSEQTGGKLPLMVTENGSAWYDP.PHAIDGRIHD......PMR

AoBgl  341  SKLSDEERSDMRRFIEAQLDAFELKSGWVFWTWKTEGAPGWDMSDLLEAGVFPTSPDDR.
NkBgl  398  VHYYTEHLKEMLKAIHE..DGV.NVIGYT..AWSLMDNFEWLRGYSEKFGIYAVDFEDPA
HiBgl  423  VKYFNDYVRAMAAAVAE..DGC.NVRGYL..AWSLLDNFEWAEGYETRFGVTYVDYANDQ
Td2F2  373  QAFFEAHLAQLQRAL.A..AGV.PLKGYF..AWSLLDNFEWAMGLSMRYGICYTNFETLE
Bgl6   382  VHYLQTHIKALHDAI.G..KGV.DLRGYM..AWSLLDNLEWSLGYSKRFGIVHVNFATQE

AoBgl  400  ..EFPKQC..............................
NkBgl  453  RPRIPKESAKVLAEIMNTRKIPERFRDLEHHHHHH...
HiBgl  478  K.RYPKKSAKSLKPLFDSLIRKE...............
Td2F2  427  R.RIKDSG.....YWLRDFIAGQRGKLAALEHHHHHH.
Bgl6   436  R.TIKDSG.....LLYAEVIKTHGDVLNTLKLHHHHHH
```

**Figure S2: Multiple sequence alignment of AoBgl with other well characterized GH1 family β-glucosidases.** The β-glucosidases considered here are from *Neotermes koshunensis* (NkBgl), *Humicola insolens* (HiBgl), and soil metagenome (Td2F2 and Bgl6).

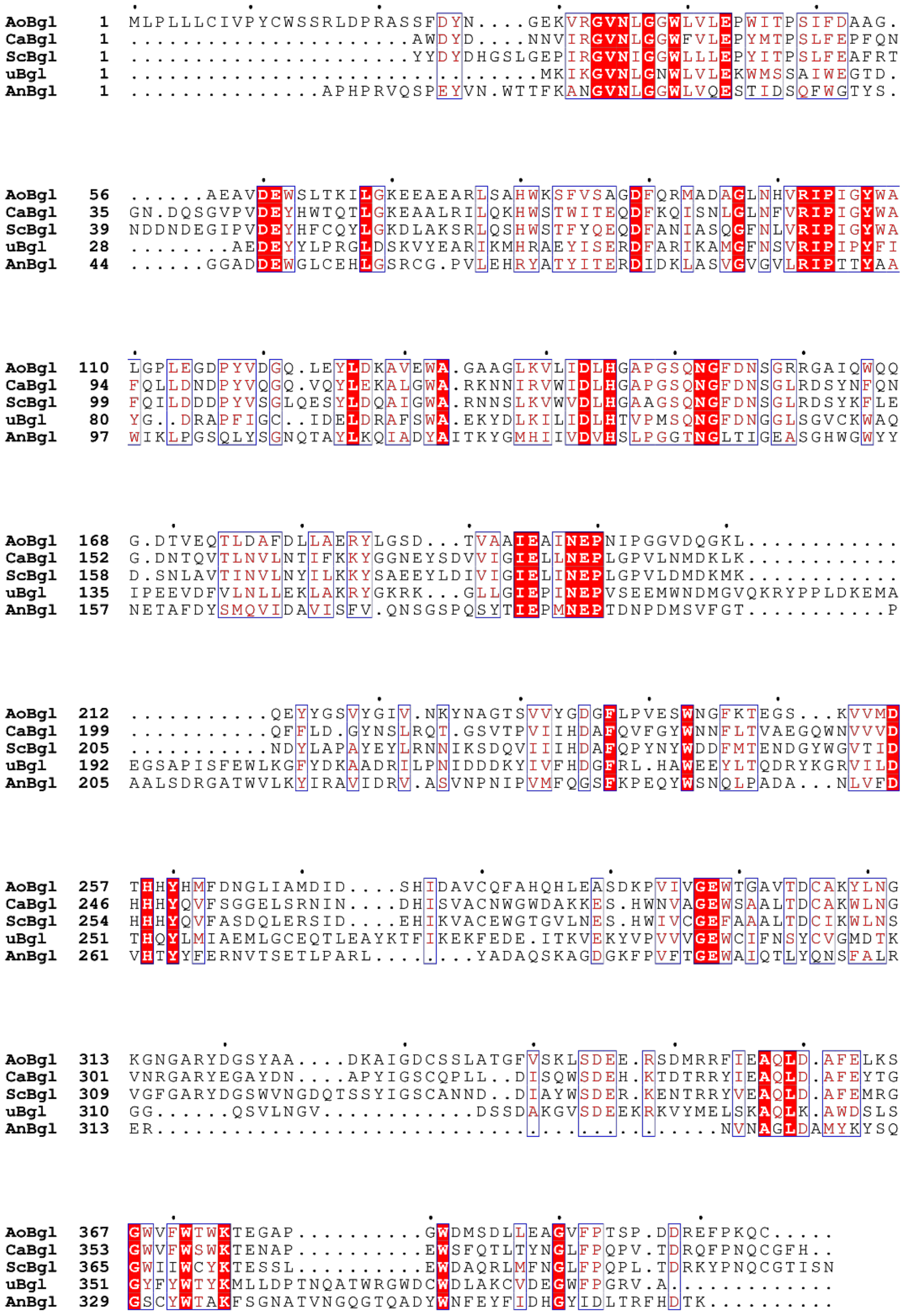

**Figure S3: Multiple sequence alignment of AoBgl with other well-characterized GH5 exo β (1,3) glucanases.** The other GH5 enzymes are from *Candida albicans* (CaBgl), *Saccharomyces cerevisiae* (ScBgl), uncultured bacterium (uBgl) and *Aspergillus niger* (AnBgl).

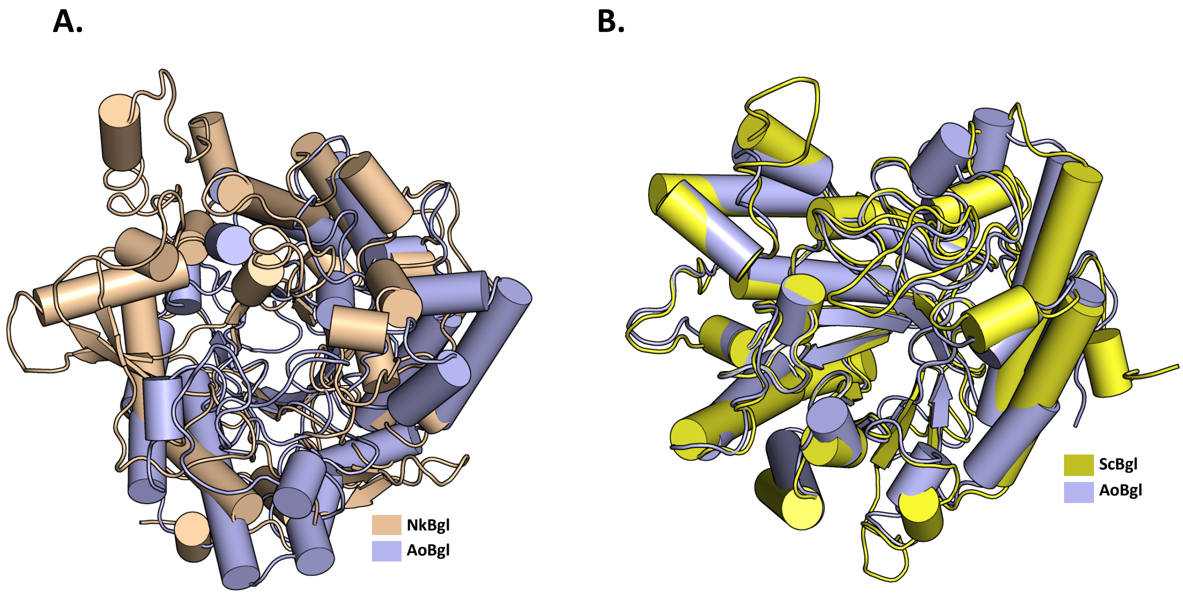


**Figure S4: Structural comparison of AoBgl with other characterized enzymes.** (A.) GH1 enzyme from *Neotermes koshunensis* (NkBgl) (PDB ID: 3VIK) and (B.) GH5 enzyme from *Saccharomyces cerevisiae* (ScBgl) (PDB ID: 1H4P).

**Supporting Data 1:**

***Experimental procedure:***

The structural stability of AoBgl at different temperatures was assessed by studying the tryptophan fluorescence. The 3µM of the purified enzyme was incubated at temperatures of 30°C, 40°C, 50°C, 55°C and 60°Cfor 1 hour and the emission spectra was recorded from 310-450 nm after excitation at 290 nm after every 5 minutes. The protein showed an emission maximum at 337 nm at 30°C which was considered as the native protein and the fluorescence intensity at this wavelength was plotted for each temperature using Graph Pad Priam 8.4.2.

***Results:***

AoBgl lost activity during incubation at 55°C, which was the calculated optimum hydrolysis temperature within 1 hour of incubation. The temperature-dependent tryptophan fluorescence was recorded to estimate the probable causes of this behaviour. The fluorescence signal at 337 nm. It was noted that during this incubation period, there was very little change in the fluorescence intensity at 337 nm at lower temperatures, however at higher temperatures of 55˚C and 60˚C, there was a steep decline in the fluorescence intensity at 337 nm (Figure S5) which indicated unfolding of the protein at these temperatures thereby exposing the buried tryptophan residues towards the aqueous environment. The unfolding of AoBgl at these temperatures might cause the observed loss of activity.

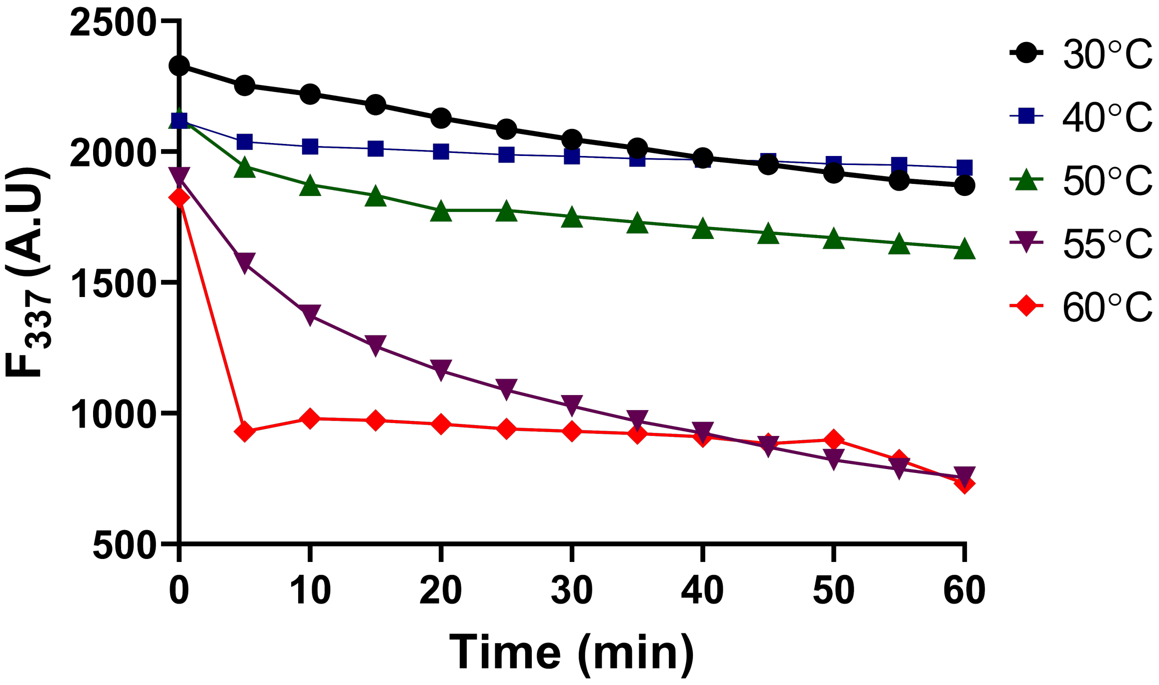


**Figure S5: Thermal unfolding in AoBgl.** Change in fluorescence emission intensity of AoBgl at 337 nm with change in temperature from 30°C to 60°C indicates that AoBgl undergoes temperature induced denaturation.

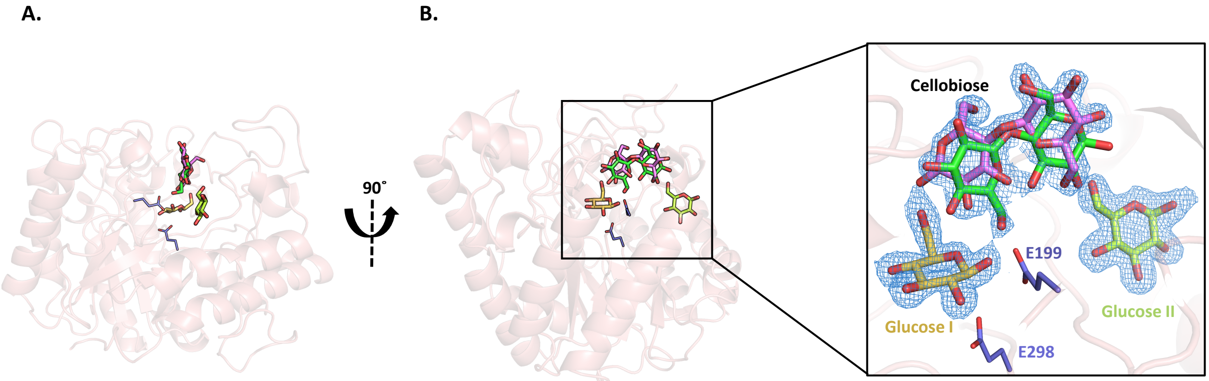


**Figure S6: Structural overview of the cellobiose-bound form of AoBgl:** (A.) The overall structure of AoBgl is shown as a salmon cartoon. (B.) 90° rotated view of the same. Catalytic glutamates (E199 and E298) are shown as blue sticks; glucose molecules are shown as dark yellow and yellow-green sticks; cellobiose in double conformation is shown as light purple and green sticks. Inset: zoomed-in view of the active site showing the bound sugars. Corresponding 2 $F_O$-$F_C$ electron densities contoured at 1σ level are shown as a sky-blue mesh. Glucose captured at active site is denoted as "Glucose I"; whereas the second glucose molecule observed towards the opening of the catalytic crater is denoted as "Glucose II".

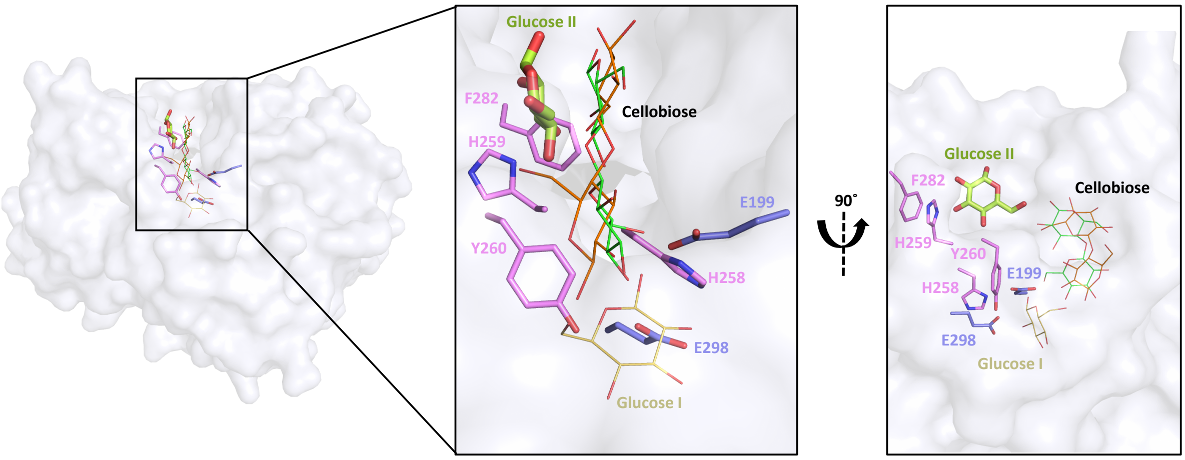


**Figure S7: Position of second glucose molecule in the cellobiose-bound AoBgl structure.** Inset: zoomed-in view of the catalytic crater and 90° rotated view of the same. The bound glucose at the catalytic site is indicated as "Glucose I" and shown as dark yellow sticks, bound cellobiose in dual conformation is shown as yellow and green sticks. The second glucose molecule captured at the crater's entrance is labelled "Glucose II" and shown as light green thick sticks. The catalytic glutamates are shown as blue sticks and other residues within 4Å of "Glucose II" are shown as purple sticks.

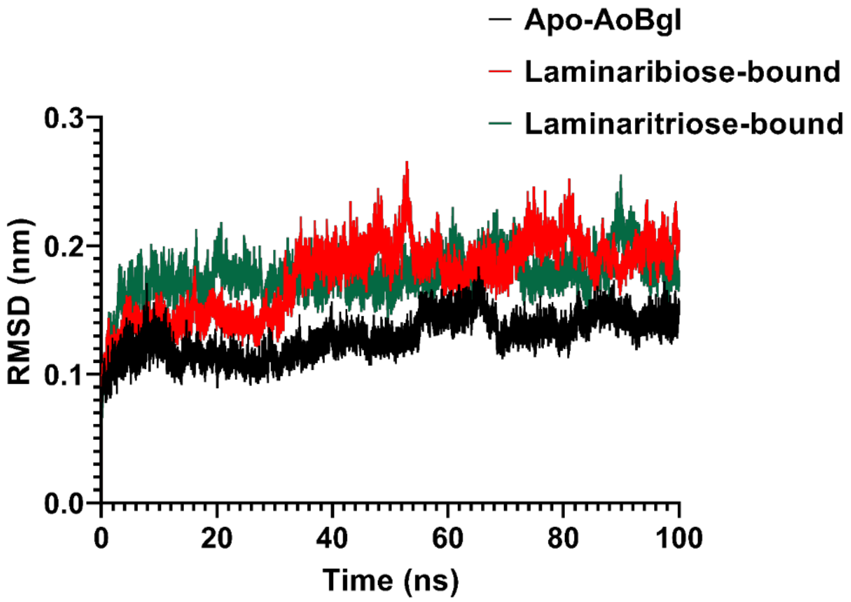


**Figure S8: Root-mean-squared deviation (RMSD) plot for the trajectories AoBgl during 100 ns of MD simulation run.** Apo-AoBgl is indicated in black, laminaribiose-bound AoBgl is indicated in red and laminaritriose-bound AoBgl is indicated in green.

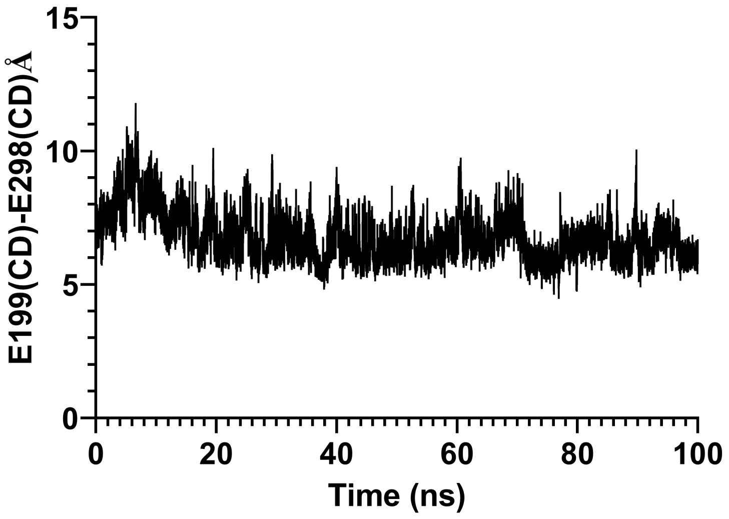


**Figure S9: Distance between the CD of the catalytic glutamates E199 and E298 during 100 ns of MD simulation run.** The distance plot shows that the two catalytic glutamates are ~5.5-6.0 Å apart, which is important for the retention mechanism of carbohydrate hydrolysis.

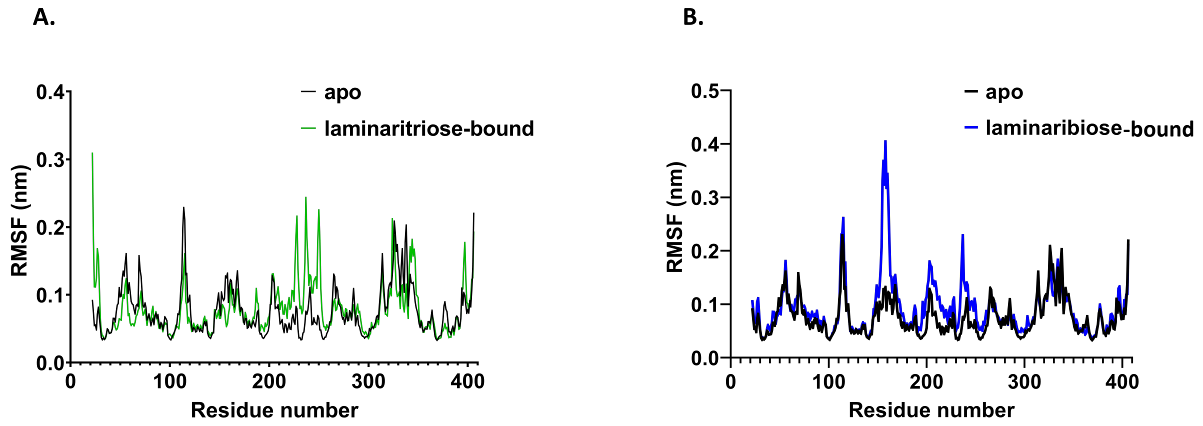


**Figure S10: Stability of AoBgl complexed with different ligands.** (A.) Comparison of RMSF plots of AoBgl in apo (black) against laminaritriose-bound form (green) and (B.) Comparison of RMSF plots of AoBgl in apo (black) against laminaribiose-bound form (blue).

**Table S1: Sequence of primers for site-directed mutagenesis in AoBgl:**

| Name of mutant | Forward primer | Reverse primer |
|---|---|---|
| AoBgl-F263A | 5′ TACCATATGGCTGACAACGGACTCATTGCCATGGATATC 3′ | 5′ GAGTCCGTTGTCAGCCATATGGTAATGGTGAGTGTCCATCAC 3′ |
| AoBgl-E298S | 5′ GTTATTGTGGGTTCGTGGACCGGTGCCGTGACTGAC 3′ | 5′ GGCACCGGTCCACGAACCCACAATAACAGGCTTGTC 3′ |

**Movies included:**

**Supplementary movie M1:** Structural characterization of AoBgl showing the active site in apo form and the interactions of the bound sugars in cellobiose-bound form. The sugar moieties in laminaritriose are found to occupy the same sites as the glucose moieties in the cellobiose-bound structure of AoBgl.

**Supplementary movie M2:** Trajectory of AoBgl with bound laminaritriose. The ligand laminaritriose remains bound inside the catalytic crater throughout the duration of MD simulation.

**Supplementary movie M3:** Trajectory of AoBgl with bound laminaribiose. The ligand laminaribiose escapes the catalytic crater within 20-40 ns of MD simulations and moves freely within the solvation box.
